# Bioorthogonal cyclopropenones for investigating RNA structure

**DOI:** 10.1101/2024.10.22.619649

**Authors:** Sharon Chen, Christopher D. Sibley, Brandon Latifi, Sumirtha Balaratnam, Robert S. Dorn, Andrej Lupták, John S. Schneekloth, Jennifer A. Prescher

**Affiliations:** Departments of Chemistry, University of California, Irvine, California 92697, United States; Molecular Biology & Biochemistry, University of California, Irvine, California 92697, United States; Pharmaceutical Sciences, University of California, Irvine, California 92697, United States; Chemical Biology Laboratory, National Cancer Institute, Frederick, Maryland 21702

## Abstract

RNA sequences encode secondary and tertiary structures that impact protein production and other cellular processes. Misfolded RNAs can also potentiate disease, but the complete picture is lacking. To establish more comprehensive and accurate RNA structure-function relationships, new methods are needed to interrogate RNA and trap native conformations in cellular environments. Existing tools primarily rely on electrophiles that are constitutively “on” or triggered by UV light, often resulting in high background reactivity. We developed an alternative, chemically triggered approach to crosslink RNAs using bioorthogonal cyclopropenones (CpOs). These reagents selectively react with phosphines to provide ketenes—electrophiles that can trap neighboring nucleophiles to forge covalent crosslinks. As proof-of-concept, we synthesized a panel of CpOs and appended them to thiazole orange (TO-1). The TO-1 conjugates bound selectively to a model RNA aptamer (Mango) with nanomolar affinity, confirmed by fluorescence turn-on. After phosphine administration, covalent crosslinks were formed between the CpO probes and RNA. The degree of crosslinking was both time and dose-dependent. We further applied the chemically triggered tools to model RNAs in biologically relevant conditions. Collectively, this work expands the toolkit of probes for studying RNA and its native conformations.

## Introduction

RNA is best known for its role as an information carrier in protein synthesis.^1^ However, only 2% of known cellular RNAs are translated. The remaining 98% comprise noncoding RNAs (ncRNAs) that play key roles in cellular processes, such as modifying chromatin and regulating mRNA abundance.^2,3^ These functions are dependent on the structural features of a given RNA, which are not static. RNA can further adopt a variety of conformations in response to various stimuli,^4–6^ and each motif can regulate a distinct biological process. For example, the G-quadruplex GQ-2 in the Bcl-x pre-mRNA has been shown to regulate apoptosis. GQ-2 is a highly structured element, and its stabilization by proteins or small molecules can trigger splicing, resulting in cell death.^7^ Although the role of GQ-2 in the Bcl-x pre-mRNA has been elucidated, the structure-function relationships of most RNAs remain largely unknown.

Teasing apart the subtle dynamic changes associated with RNAs (and their downstream effects) requires methods to examine RNA structural heterogeneity in physiologically relevant environments.^8–12^ Since sequence information alone cannot be used to accurately predict dynamics, methods to examine the full scope of structural variability are needed. Classic biophysical approaches (e.g. X-ray crystallography^13–16^ and NMR^17–19^) offer high structural resolution, but place limitations on RNA size and environment. Chemical probing methods, by contrast, can tag RNAs of all lengths in live cells.^20–22^ These reagents selectively label solvent accessible nucleobases^23–26^ or 2’-hydroxyl groups, and when combined with next generation sequencing, they can be used to predict base-pairing interactions associated with RNA secondary structure^27–30^. However, most chemical probes are “always on” and cannot be used to investigate RNAs in response to discrete stimuli. They are thus limited in their ability to report on conformational dynamics. Photoactivatable versions have been developed to address this issue, but cannot be easily translated to more heterogeneous environments owing to their need for UV irradiation.^31^

To address the need for more biofriendly RNA probes, we aimed to develop bioorthogonal chemistries for examining RNA structures. Many bioorthogonal reagents are compact, imparting minimal-to-no perturbation to the target biomolecule. In addition, bioorthogonal groups can react on demand and enable capture of motifs. One such handle that is of particular interest to our groups is the cyclopropenone (CpO). CpO motifs are small functional groups that react with biocompatible phosphines to generate electrophilic ketene-ylide intermediates (Figure 1). Neighboring nucleophiles (e.g., hydroxyl groups and nucleobases) can then trap the ketene, leading to a crosslinked product. One can initiate reactivity at any time, enabling RNA structural changes to be examined in response to various stimuli.

**Figure 1.**
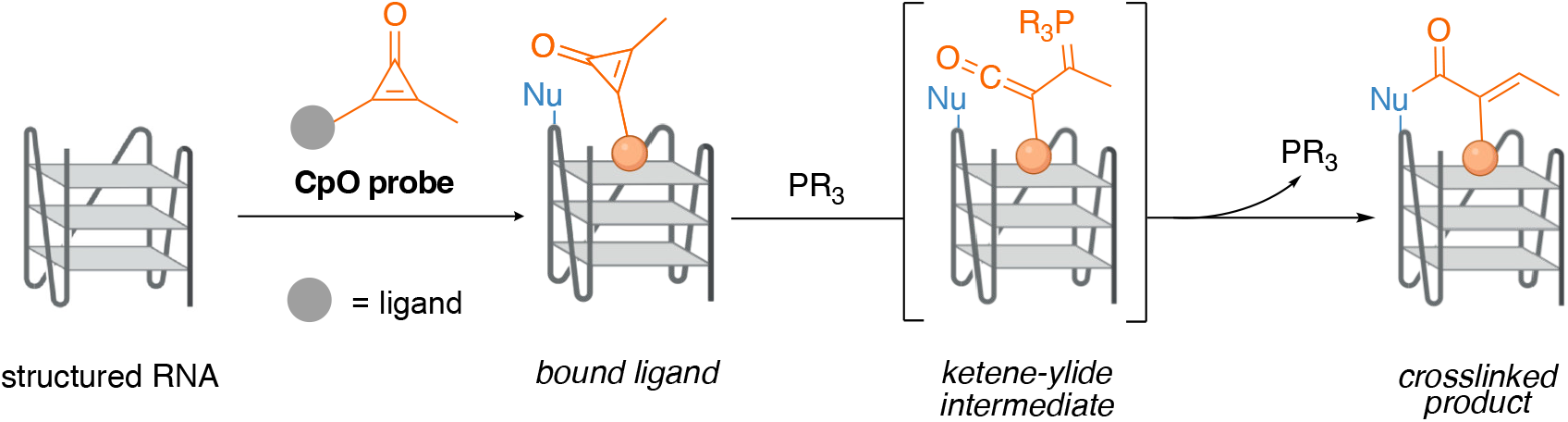
Cyclopropenones as chemically triggered probes for examining RNA structure. In this work, the Mango aptamer was used as a model to demonstrate CpO-mediated crosslinking.

Here, we report a platform for sampling RNA structures using bioorthogonal CpO motifs. We previously applied the CpO-phosphine reaction to both label and crosslink proteins.^32,33^ In this work, we show that the bioorthogonal reaction can be used similarly to study RNA conformations. The reaction was evaluated with model nucleophiles for trapping structured RNA motifs. Crosslinking was further examined with a model aptamer-ligand pair. Collectively, this work establishes a new chemical method for examining RNA structure using a conditionally activated chemical probe.

### Design, synthesis and characterization of TO-1-CpO

To develop a bioorthogonal platform for triggered crosslinking, we initially focused on the fluorogenic RNA aptamer Mango (Figure 1). Mango is a laboratory-evolved, highly structured RNA that forms a three-tiered G-quadruplex motif and binds the fluorogenic dye thiazole orange biotin (TO-1-biotin) with low nanomolar affinity.^34^ This ligand is synthetically accessible and exhibits enhanced fluorescence turn-on upon binding to Mango.^35–37^ Unrau and colleagues reported that two polyethylene glycol PEG units inserted between the TO-1 and biotin motifs provided the lowest *K*_D_ values and highest fluorescence turn-on.^34^ We thus aimed to construct a related TO-1 variant comprising a bioorthogonal CpO motif (**TO-1-CpO**, Scheme 1). The synthesis involved an initial conjugation of CpO-PFP to commercially available *t*-Boc-*N*-amido-PEG_2_-amine, followed by acid deprotection.^32^ The CpO-PEG_2_-amine product was then coupled with TO-1 acetate to provide the desired probe.^38^

**Scheme 1.**
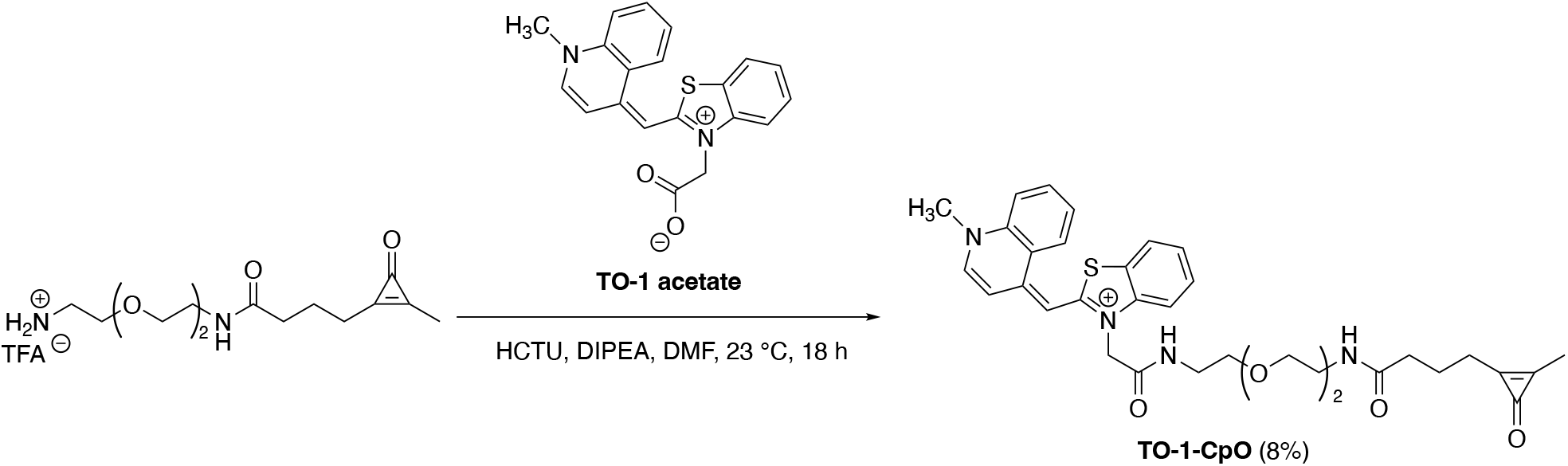
Synthesis of TO-1-CpO.

With **TO-1-CpO** in hand, we first examined whether the probe could be activated by bioorthogonal phosphines and subsequently trapped. Phosphine treatment was expected to initiate a crosslink between the ligand and the aptamer only in the presence of nearby nucleophiles. The absence of a suitable trap would instead yield an innocuous hydrolyzed product (Figure 2A). We also surmised that ribose hydroxyl groups could serve as nucleophiles, based on earlier work demonstrating that tyrosine phenol groups could forge crosslinks with CpO-modified proteins.^32^ We first simulated potential nucleotide trapping with aniline and benzylamine. **TO-1-CpO** was incubated with the amine traps in the presence of PTA, a commercially available and water-soluble phosphine used in previous studies.^39^ The reactions were monitored by HPLC, and crosslinked products were observed in the presence of either excess benzylamine or aniline (Figures 2B-C). In the absence of any additional trapping reagent, hydrolyzed (quenched) products were formed. Mass spectrometry was used to confirm the identities of the species (Figure S1). Collectively, these results suggested that nucleophiles present on nucleotides could react with activated CpO motifs.

**Figure 2.**
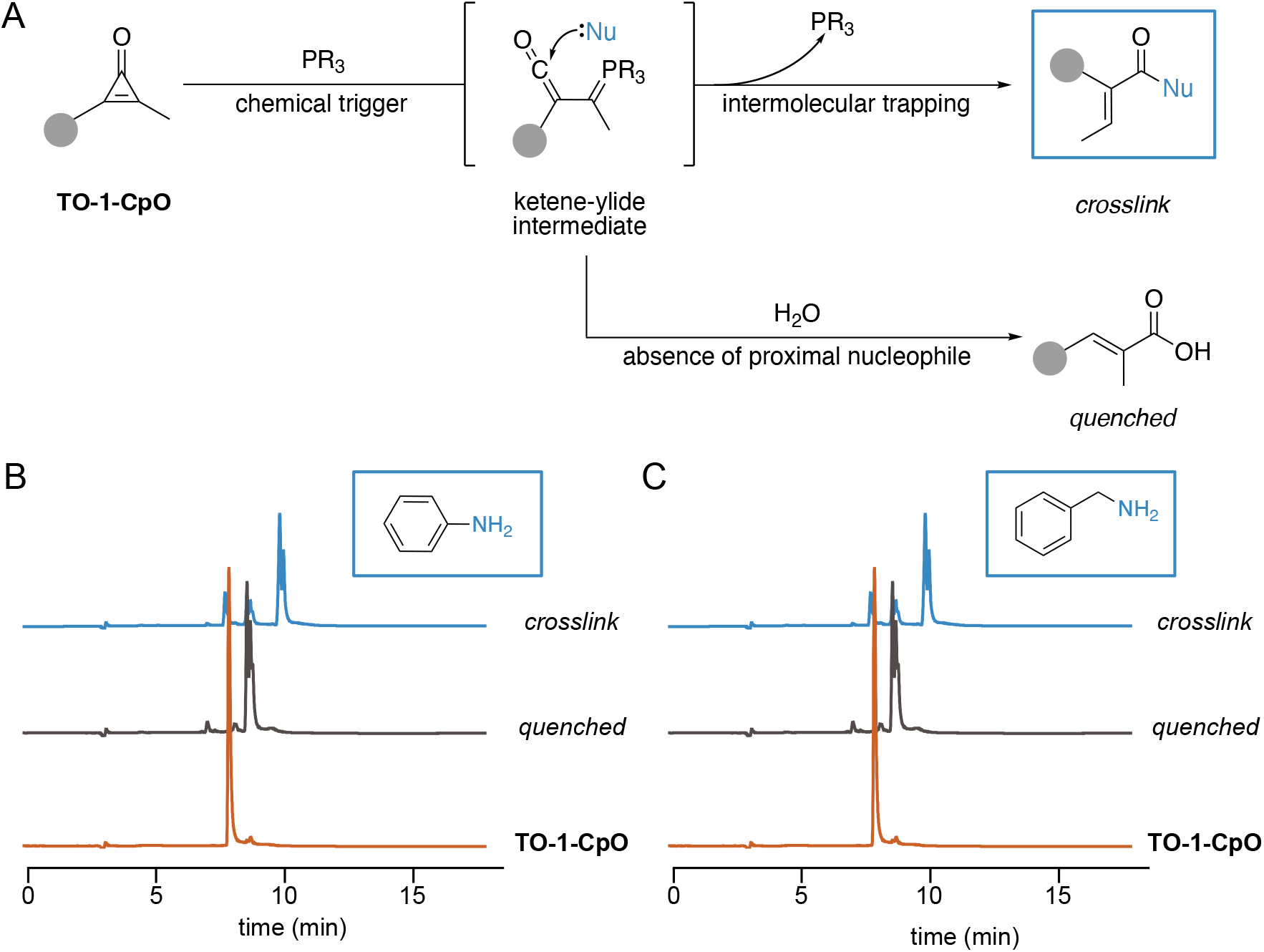
TO-1-CpO is activated upon phosphine administration and trapped by nucleophiles. (A) Overall design of *in vitro* trapping experiments. (B-C) HPLC traces of **TO-1-CpO** and the quenched and crosslinked products. **TO-1-CpO** was incubated with PTA without any nucleophile to determine the retention time of hydrolyzed product. Ketene-ylide intermediates were trapped with (B) aniline or (C) benzylamine to forge covalent linkages.

### Investigating the binding and crosslinking between TO-1-CpO and Mango

We next pursued CpO-mediated crosslinking with the Mango aptamer as a model RNA. Given the four distinct variants of Mango availble,^40–43^ each with its own unique structure, we initially examined the ligand-bound crystal structures (Figure 3A). The native biotin fragment was used to approximate the location of CpO with respect to the aptamer. The models suggested that the CpO unit would remain solvent exposed in Mango II, and thus accessible to the phosphine trigger. We therefore moved forward with this aptamer. **TO-1-CpO** was incubated with Mango II, and fluorescence measurements were used to confirm ligand binding. As expected, signal turn-on was comparable to that observed with known ligand TO-1-biotin (Figure 3B). The binding affinities (*K*_D_) for the remaining Mango aptamers were also determined. **TO-1-CpO** exhibited the tightest binding with Mango II. Importantly, binding was dramatically reduced with an inactive Mango II mutant, where the guanines involved in G-quadruplex formation were mutated to cytidines (Figure 3C).

**Figure 3.**
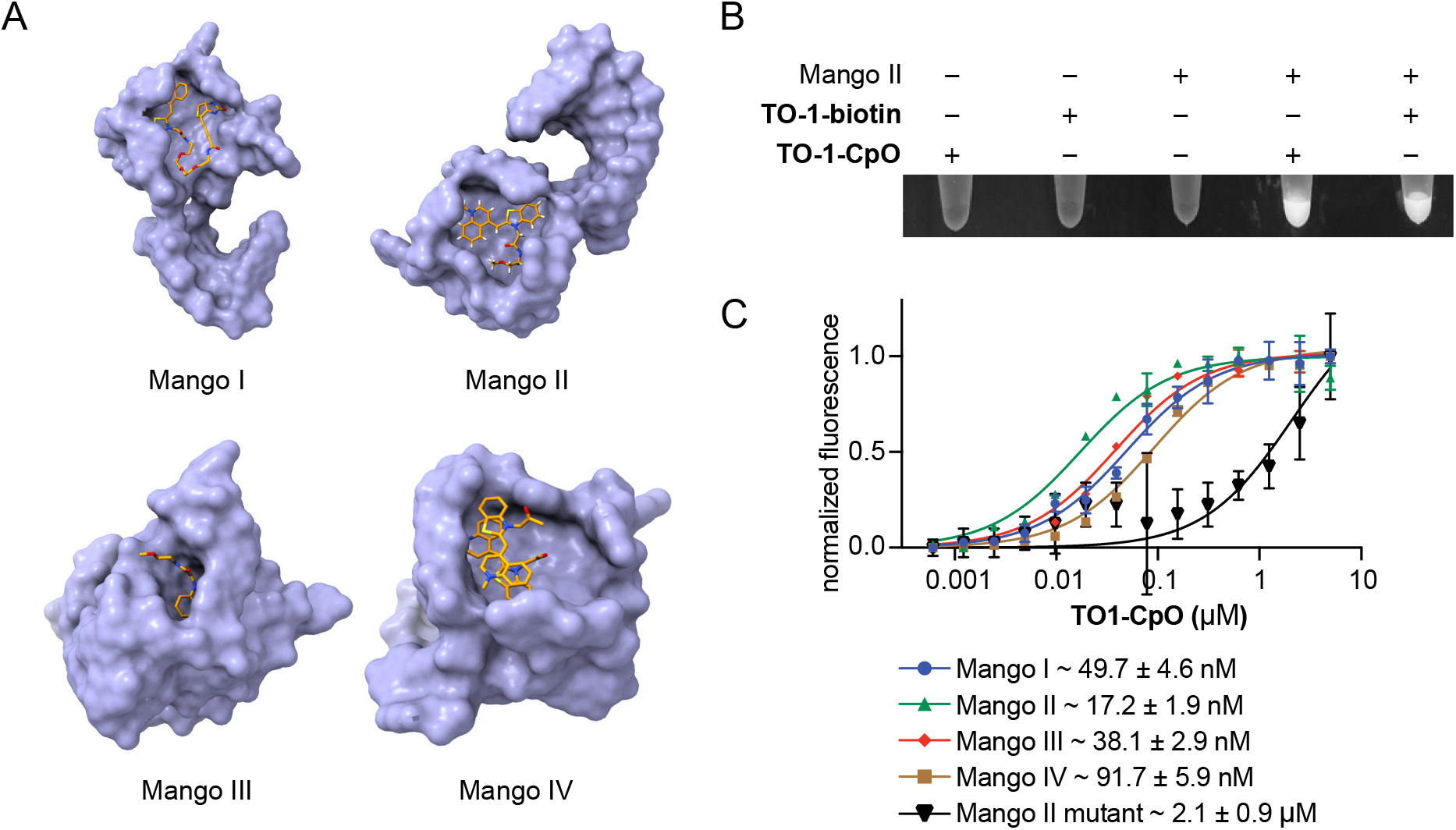
TO-1-CpO binds Mango II with nanomolar affinity. (A) Crystal structures of each Mango aptamer with TO-1-biotin depicted as space filling models. Mango I (PDB: 5V3F), Mango II (PDB: 6C63), Mango III (PDB: 6E8S), Mango IV (PDB: 6V9D). (B) Fluorescence turn-on analyses of Mango II samples incubated with TO-1-biotin, **TO-1-CpO**, or no reagent. (C) Fluorescence measurements resulting from **TO-1-CpO** (0–10 µM) incubated with each Mango aptamer. These data were used to calculate binding affinity (*K*_D_) of **TO-1-CpO** for each flavor of Mango, reported below the graph. Error bars represent the standard error of the mean for *n* = 3 experiments.

Once ligand binding was confirmed, we aimed to crosslink **TO-1-CpO** to the Mango II aptamer and examine the effect of **TO-1-CpO** concentration on crosslinking efficiency. 5′-Cy5-labeled Mango II was incubated with varying amounts of **TO-1-CpO** (5–1000 µM) and the crosslinking reactions were assessed with denaturing PAGE. Gels were scanned using differential filters to visualize both the unmodified and crosslinked RNAs. Crosslinking was first observed with 10 μM ligand, and the fluorescence intensity of the band under the TO filter increased with larger amounts of ligand present (Figure 4B). All subsequent experiments were performed with excess ligand to maximize crosslinking yield.

**Figure 4.**
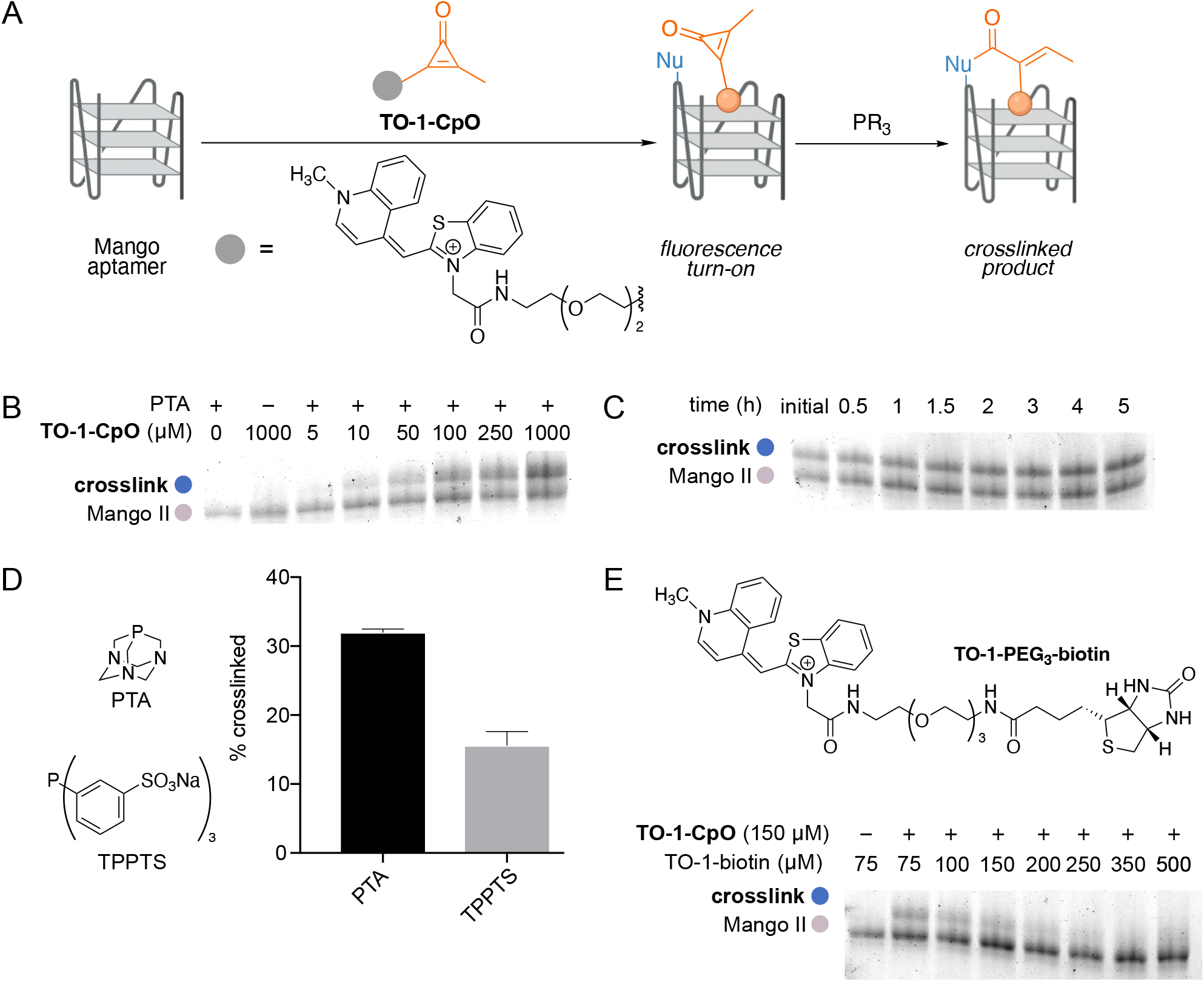
CpO-mediated crosslinking of Mango II is dependent on ligand concentration, time, and phosphine nucleophilicity. (A) Overall scheme of **TO-1-CpO** crosslinking. The ligand was incubated with Mango aptamers for 10 min, prior to phosphine addition. (B) Denaturing PAGE analysis of crosslinking reactions. 5′-Cy5 labeled Mango II (10 μM) was incubated with varying concentrations of **TO-1-CpO** and PTA (10 mM). (C) Denaturing PAGE analysis of crosslinking time dependence. 5′-Cy5 labeled Mango II (10 μM) was incubated with **TO-1-CpO** (250 μM) and PTA (10 mM), and samples were analyzed over 5 h. (D) Quantification of relative crosslinking yields with various phosphine triggers. 5′-Cy5 labeled Mango II (10 μM) was incubated for 2 h with various phosphines (10 mM), and reaction products were analyzed via denaturing PAGE. Error bars represent the standard error of the mean for independent replicate experiments (*n* = 4 for PTA and n = 3 for TPTPS). (E) Denaturing PAGE analysis of crosslinking experiments performed in the presence of competing ligand (TO-1-biotin). TO-1-biotin (75–500 μM) was added to reactions comprising Mango II (10 μM), **TO-1-CpO** (150 μM) and PTA (10 mM).

### Optimizing CpO-mediated crosslinking conditions

We further evaluated the time and temperature dependence of the reaction to maximize crosslinking yields. Crosslinking increased over two hours prior to reaching a plateau (Figure 4C). Increased labeling was also observed at higher temperatures (Figure S3). Although crosslinking yields at 25 °C and 37 °C were similar, 37 °C was chosen for subsequent experiments to mimic physiological conditions.

We next examined the effect of phosphine nucleophilicity on relative crosslinking yield. More reactive phosphines enable rapid crosslinking, but these same reagents are more susceptible to oxidation.^32^ Ten phosphines were examined using the optimized parameters (Table S1). Included in this group were well known phosphine ligands (TPP, CyDPP, PTA, and PTABS) and biological reagents (THMP, THPP, and TCEP). Crosslinking was evaluated using denaturing PAGE as above, and relative efficiencies were calculated by gel densitometry. Differential crosslinking was observed across the panel, and the best performing phosphines generally included the most nucleophilic reagents (TPP, CyDPP, PTA, THMP). We did not move forward with TPP, CyDPP, or THMP, though, given the poor solubility of these reagents in aqueous buffers. In addition, the negatively charged phosphine reagents (TPPTS and CyDPPDS) resulted in less crosslinking compared to their neutral counterparts. We attributed this result to charge repulsion with the phosphate backbone of RNA. Based on the remaining phosphines that exhibited good labeling yields, PTA was chosen over PTABS to minimize additional charge repulsion effects (Figure 4D).

### Examining the specificity of CpO-mediated crosslinking

We further investigated whether **TO-1-CpO** binding to the target RNA was necessary, or if crosslinking was ubiquitous regardless of the aptamer present. In one experiment, we included Mango’s cognate ligand, TO-1-biotin, as a competitive binder to the aptamer. Reactions were performed with **TO-1-CpO** held at a constant concentration (150 μM) and varying TO-1-biotin (75–500 μM). Crosslinking decreased as concentration of the competitor increased, suggesting that both ligands occupy the same aptamer binding site (Figure 4E). We also examined additional probes lacking key structural elements for crosslinking, TO-1-NH_2_ and Ph-CpO. TO-1-NH_2_ contains the dye and can bind to the Mango II aptamer, but lacks the CpO moiety, whereas Ph-CpO cannot localize to the aptamer, but contains the CpO motif. Importantly, when these negative control probes were used in place of **TO-1-CpO**, crosslinking was not observed. These observations confirm that each structural element of **TO-1-CpO** plays an essential role in crosslinking to the RNA (Figure S4A).

The specificity of the labeling reaction was further examined using other structured nucleic acids. The panel included a range of motifs [e.g., hTERT (an RNA pseudoknot), cMyc IRES (an RNA hairpin), or RNAse P (a complex RNA)] not known to interact with TO-1. As expected, minimal to no-crosslinking was observed when non-target RNAs were incubated with **TO-1-CpO** and PTA (Figure S4B-C). Specificity for Mango II was further showcased when **TO-1-CpO** was incubated with an inactive variant of the Mango II aptamer, where the guanines involved in forming the ligand-binding G-quadruplex structure were mutated to cytidines (Figure S5). Minimal crosslinking was observed at ligand concentrations below 100 μM. Some covalent trapping was observed at higher concentrations, but the efficiency was much lower than that observed with wild-type Mango II.

**TO-1-CpO** was further incubated with other Mango aptamers in the presence of phosphine. Crosslinked product formation was analyzed by denaturing PAGE and MALDI-MS (Figure S6). The most labeling was achieved with Mango II, confirming our hypothesis regarding accessibility (Figure S7). Crosslinking with other Mango aptamers was also observed, but to a lesser extent. Collectively, these results demonstrate that CpO-mediated crosslinking between **TO-1-CpO** and Mango is specific to both the RNA sequence and three-dimensional structure.

### Identifying the proximity dependence and location of crosslink

To gain insight into the specificity of crosslinking, we incubated 5′ Cy5 labeled Mango II, 3′ Cy5 Mango II, and TO-1-CpO crosslinked Mango II with RNAse T1. This nuclease cleaves specifically after guanines, which are highly abundant in the Mango aptamer. The resulting cleavage patterns were evaluated using denaturing PAGE (Figure S8). A laddering effect was observed with the 5′ Cy5 labeled Mango II and 3′ Cy5 labeled Mango II samples. The same pattern was not observed with **TO-1-CpO** crosslinked aptamer, suggesting that the activated probe does not non-specifically label the termini.

To identify specific sites of crosslinking, 5′ Cy5 labeled Mango II and the **TO-1-CpO** crosslinked variant were reverse transcribed with MMLV reverse transcriptase. This enzyme was chosen because of its high sensitivity to RNA modifications. In our case, the TO-1 moiety was likely large enough to terminate reverse transcription at the sites of labeling and thus reveal the positions of interest. Reverse-transcribed samples were amplified and submitted for high-throughput sequencing. Terminations at positions 17, 20, 21, 22, and 25 were significantly higher with the **TO-1-CpO** crosslinked Mango II when compared to the (uncrosslinked) 5′ Cy5 labeled Mango II (Figure 5). When these termination sites were mapped onto the crystal structure, the proximity dependence of the crosslinking was immediately apparent (Figure 5A-B). Labeled nucleotides (blue) are localized near the putative CpO position (dark pink), in the top half of the aptamer. Significant termination sites were not observed on other parts of the aptamer, demonstrating that the reaction is highly specific and dependent on **TO-1-CpO** recognizing the binding site on Mango II.

**Figure 5.**
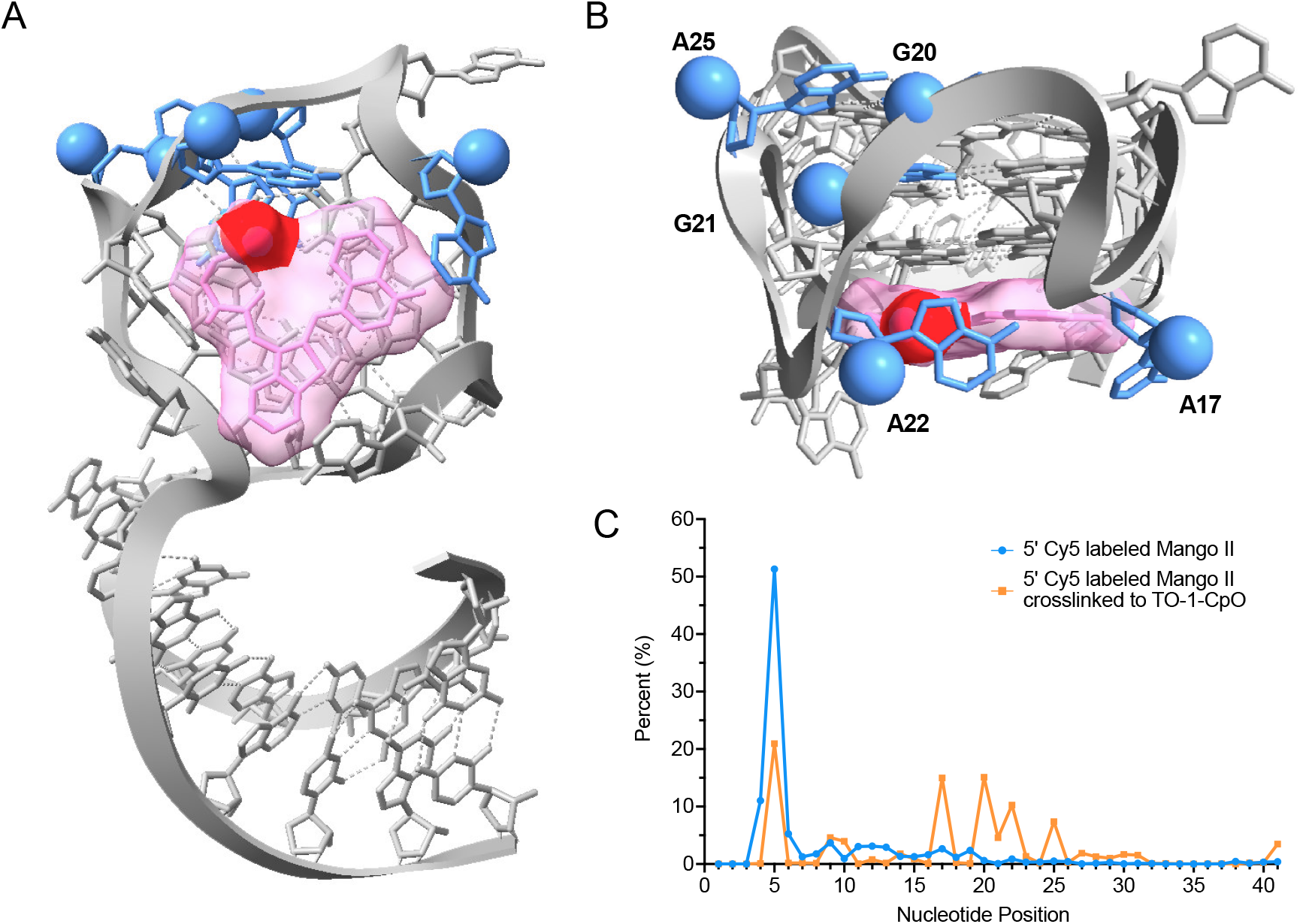
Proximity-dependent crosslinking was observed with **TO-1-CpO** and Mango II. (A) Crystal structure of Mango II aptamer docked with truncated TO-1-biotin. Approximate position of the CpO motif is highlighted in red. Sites of reactivity are colored in blue. (B) Close up of the binding domain highlighting the five modified bases. (C) Sequencing analyses of reverse transcribed Mango II aptamer. RT stops observed with TO-1-CpO modified Mango II aptamer (blue) were significantly higher at five sites in comparison to the unmodified RNA (orange).

## Conclusions

Structure plays a critical role in RNA function, and elucidating how specific conformations influence molecular recognition is an important step in understanding biological processes. Great progress has been made in deciphering RNA structure-function relationships using SHAPE and other tools.^28,44–46^ However, most of these reagents lack temporal control, making it hard to study dynamic changes. Photoactivable versions have been developed to address this issue, but the necessary irradiation is potentially harmful to nucleic acids. The community field would benefit from biocompatible and conditional (“triggerable”) probes to study the dynamics of RNA structure.

Here, we showcased CpO motifs for interrogating RNA structural dynamics by covalently modifying RNAs near ligand binding sites. CpOs were chemically activated by phosphines to tag three-dimensional RNA structures. Triggered crosslinking was achieved using the Mango II aptamer as a model system. We further demonstrated that the reaction is dependent on ligand concentration, time, temperature, and phosphine nucleophilicity. In addition, we determined that the labeling is specific to the probe and proximity dependent. Finally, reverse-transcription followed by sequencing revealed the aptamer positions where covalent linkages were forged. Importantly, sites of modification were observed proximal to the known ligand binding site, indicating that specific molecular recognition of the probe is important for labeling bases that are distal in sequence but proximal in three-dimensional structure.

Future directions of this work will focus on expanding the scope and efficiency of CpO-mediated crosslinking. Applications in cells and other physiologically relevant environments are envisioned, although considerations for probe delivery and phosphine accessibility must be made. Appropriately modified reagents could be used to examine RNA conformational dynamics in response to stress, protein/ligand binding, and other stimuli. Additionally, the bioorthogonal crosslinking strategy could be used to identify biomolecules interacting with RNAs of interest, to covalently label RNAs with tags, or to map binding sites of ligands to complex RNAs. In this case, designer ligands with longer linkers could bridge (and potentially trap) neighboring proteins. Such experiments will continue to expand our understanding of RNA structure-function relationships and molecular recognition of RNA in biological settings.

## Supporting Information

The accompanying file contains general materials and methods, Figures S1-S9, Table S1, synthetic procedures, and small molecule/conjugate characterization data.

## Notes

The authors declare no competing financial interest.

## Acknowledgements

This work was supported by the U.S. National Institutes of Health (R01 GM 1262226 to J.A.P.), the Chan-Zuckerberg Initiative (J.A.P.) and Intramural Program of the NIH, NCI, Center for Cancer Research (J.S.S.). We thank Benjamin Katz and Felix Grün at the UC Irvine Mass Spectrometry facility and the NCI Biophysics Resources center for assistance with mass spectrometry. We also thank members of the Prescher, Lupták, and Schneekloth labs for insightful discussions.

