## Supplementary Materials for "Bioorthogonal cyclopropenones for investigating RNA structure"

**Contents**


I. **Supplementary figures**

II. **Binding studies**

III. **Crosslinking experiments**

IV. **Elucidation of crosslinking location**

V. **Oligonucleotides**

VI. **General synthetic procedures**

VII. **Synthetic procedures**

VIII. **NMR spectra**

IX. **References**

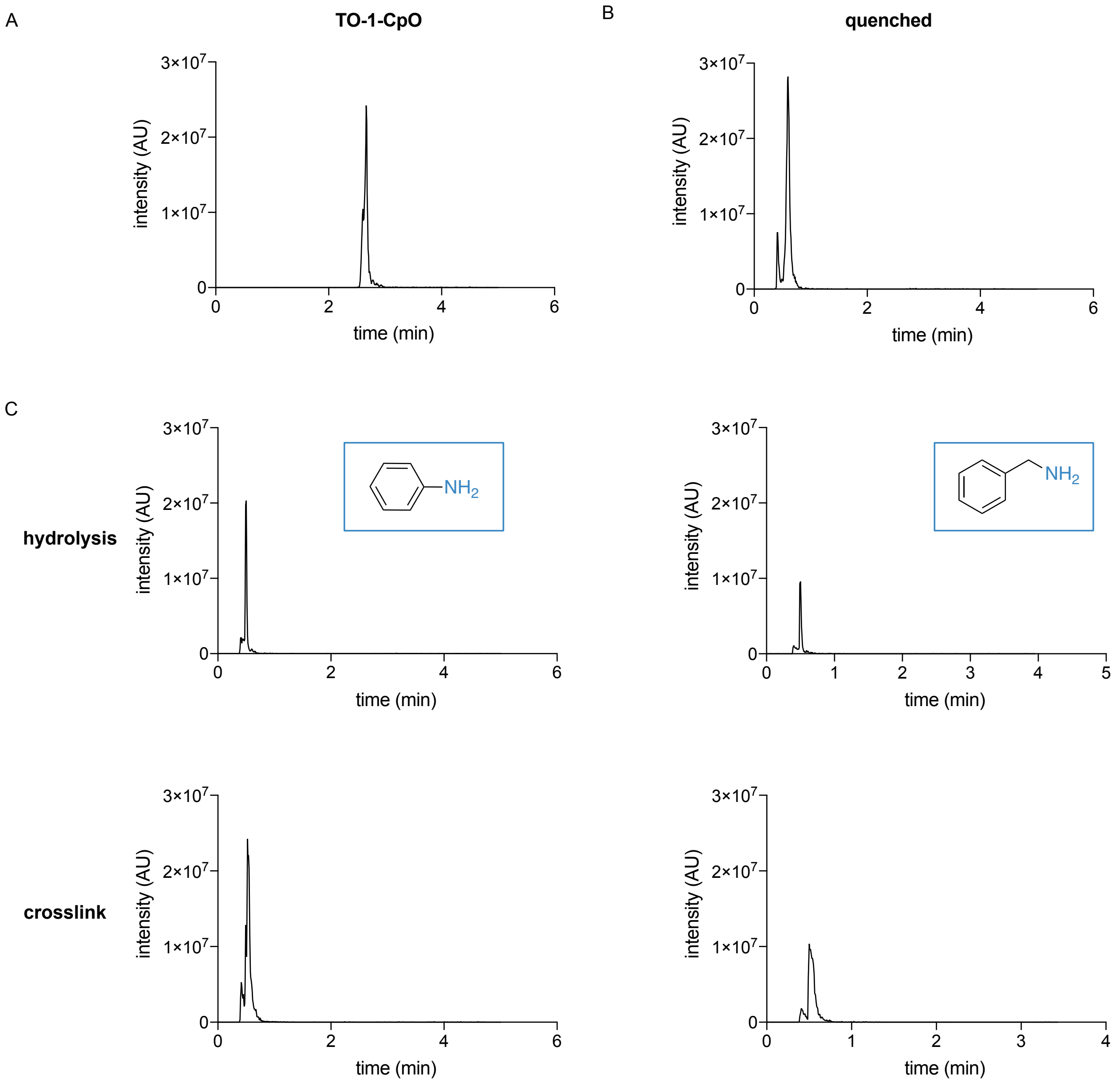


**Figure S1. Analysis of CpO hydrolysis and in vitro trapping via LC-MS**. Extracted ion chromatograms (EICs) of samples containing (A) only **TO-1-CpO** (*m/z* = 615), (B) **TO-1-CpO** and PTA, or (C) **TO-1-CpO**, PTA, and aniline. In the absence of a trapping nucleophile in (B), quenched CpO is observed (*m/z* = 633). Activated CpO trapping with (C) aniline or benzylamine provided covalent adducts (*m/z* = 708 or 722, respectively).

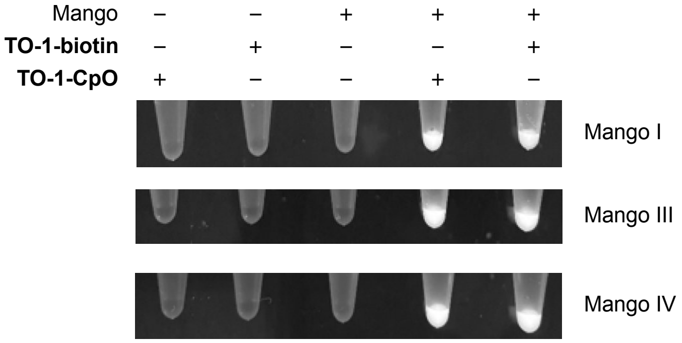


**Figure S2.** Fluorescence turn-on analysis of Mango aptamers incubated with **TO-1-biotin**, **TO-1-CpO**, or no reagent.

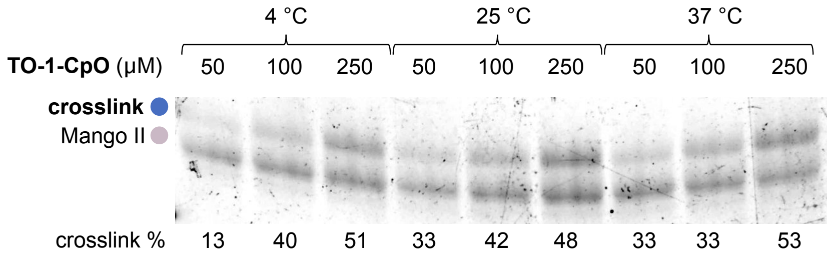


**Figure S3.** Denaturing PAGE analysis of crosslinking experiments performed with **TO-1-CpO** (50–250 µM) and PTA at varying temperatures.

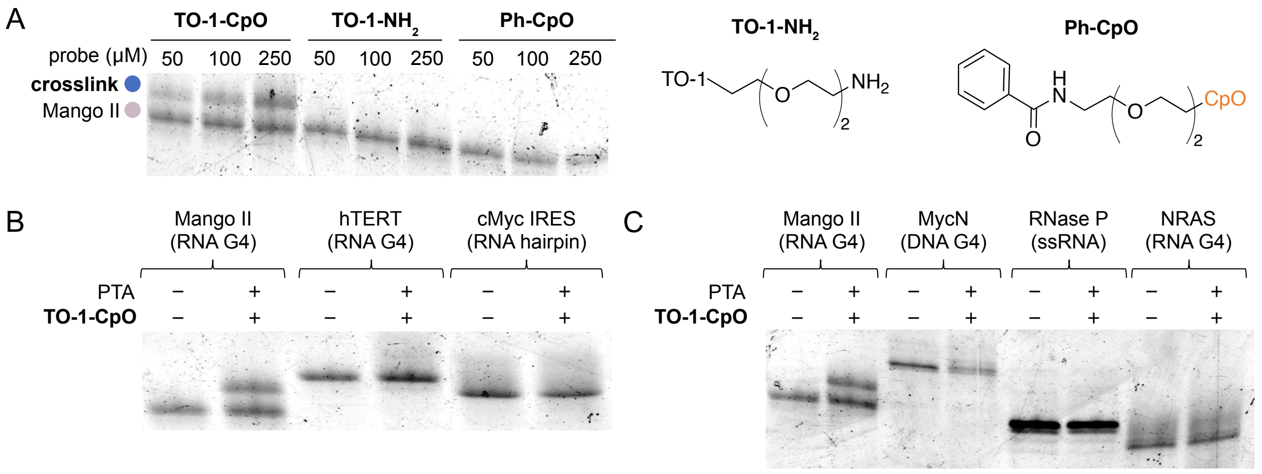


**Figure S4. CpO-mediated crosslinking is specific to the probe and aptamer**. (A) Denaturing PAGE analysis of crosslinking experiments performed with either negative control probes (**TO-1-NH$_2$**, **Ph-CpO**) or **TO-1-CpO** and phosphine. (B-C) Denaturing PAGE analysis of crosslinking experiments performed with **TO-1-CpO** and phosphine in the presence of other (non-targeted) structured RNAs.

**Table S1.** Crosslinking efficiency with a panel of commercially available phosphines.

| entry | phosphine | relative phosphine performance |
|---|---|---|
| **THMP** | 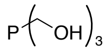 | 1.0 |
| **CyDPP** | 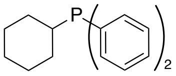 | 0.83 |
| **TPP** | 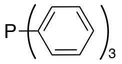 | 0.78 |
| **PTA** | 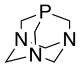 | 0.75 |
| **PTABS** | 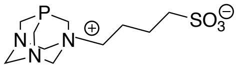 | 0.73 |
| **CyDPPDS** | 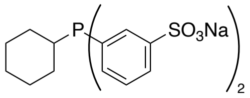 | 0.57 |
| **THPP** | 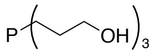 | 0.52 |
| **TPPTS** | 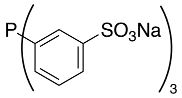 | 0.41 |
| **DCyPDIBPS** | 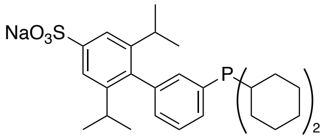 | 0.27 |
| **TCEP** | 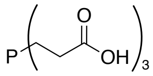 | 0.11 |

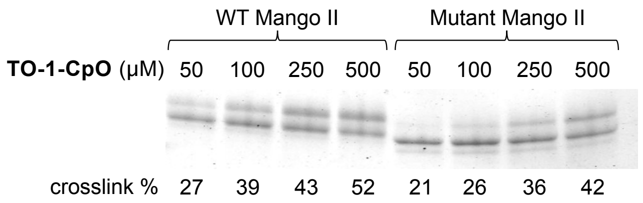


**Figure S5.** Denaturing PAGE analysis of crosslinking experiments performed with **TO-1-CpO** (50–250 μM) and phosphine in the presence of Mango **II** or a mutant aptamer.

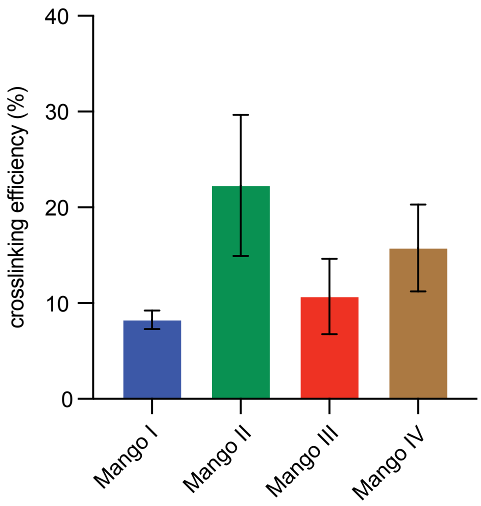


**Figure S6.** Crosslinking yields achieved with **TO-1-CpO**, phosphine, and various Mango aptamers.

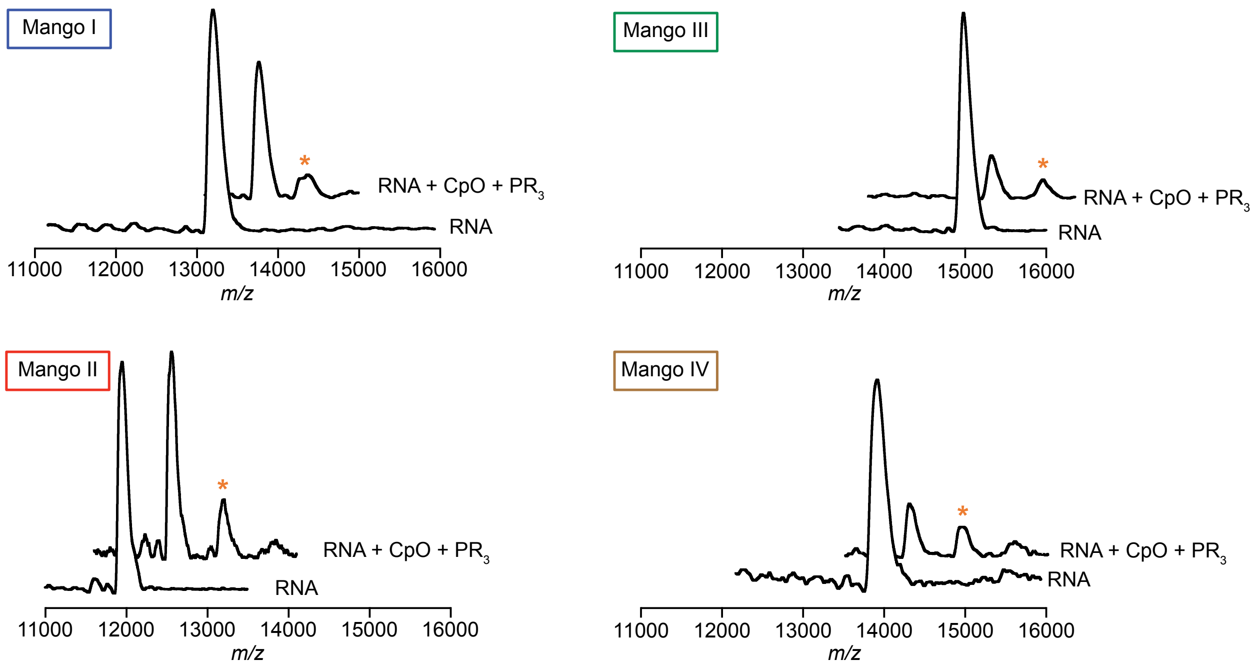


**Figure S7.** MALDI-MS spectra of crosslinking reactions performed with **TO-1-CpO**, phosphine ($PR_3$), and various Mango aptamers.

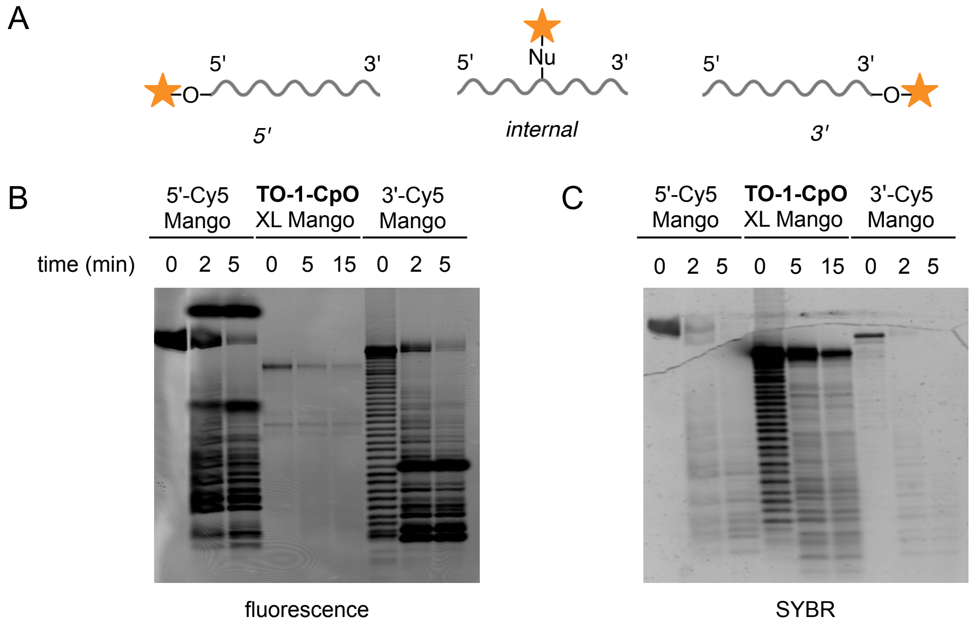


**Figure S8.** (A) Cartoon depicting potential sites of crosslinking (5' end, internal, or 3' end). (B-C) Comparison of T1 digestion patterns from 5'-Cy5 Mango, **TO-1-CpO** crosslinked Mango, and 3'-Cy5 Mango samples. The gel was visualized using both (B) Cy5 and (C) SYBR filters.

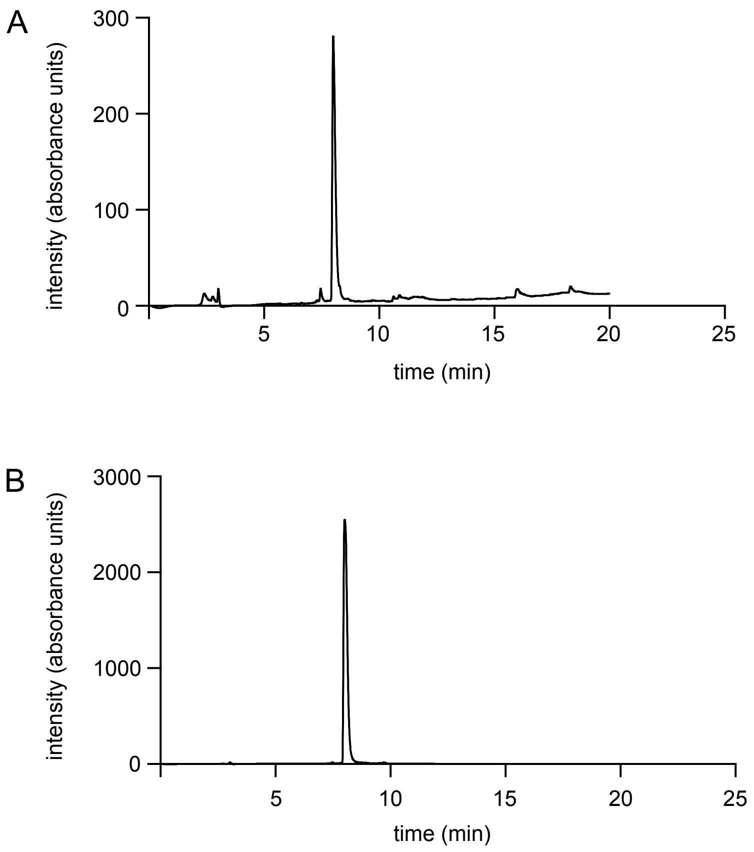


**Figure S9.** HPLC traces of purified **TO-1-CpO** monitored at (A) $\lambda$ = 254 nm and (B) $\lambda$ = 500 nm.

**Ligand-aptamer binding studies**
Fluorescence titrations were performed using unlabeled RNAs (Mango-I, Mango-II, Mango-III, Mango-IV, and Mango-II Mutant) purchased from Horizon Discovery with 10 nM **TO1-CpO** (**1**). All RNAs were prepared in buffer (10 mM Tris pH 7.5, 140 mM KCl and 1 mM $MgCl_2$) and folded by heating to 95 °C for 3 min, followed by cooling on ice for 30 min. In a black 96-well plate, **TO-1-CpO** (**1**) was prepared at a final concentration of 10 nM in 5% DMSO in buffer. After shaking for 5 min, folded RNA was added to final concentrations ranging from 0.6 nM to 5 μM. Samples equilibrated for 30 min while shaking. The fluorescence intensity was then measured on a Synergy Mx microplate reader (BioTek) at an excitation wavelength of 510 nm and emission wavelength of 535 nm. The fluorescence intensity was then normalized to the values obtained for **TO-1-CpO** (**1**) without RNA and was plotted against RNA concentration. The apparent dissociation constants were determined using a single site model to fit the curve. Each condition was performed in triplicate.

**Crosslinking experiments**

*General procedure*
Mango aptamers (10 µM, 10 µL) were resuspended in 1X folding buffer (10 mM Tris (pH 7.0), 140 mM KCl, 1 mM $MgCl_2$) and heated at 95 °C for 2 min followed by ice five minutes. The aptamer samples were then treated with TO-1 ligands (250 µM) and incubated at 37 °C with agitation. The reaction vessels were also covered with aluminum foil following ligand addition, to protect the samples form light. After 10 min, the reaction mixtures were treated with phosphine (10 mM) or no reagent, then incubated at 37 °C for an additional 2 h with shaking. The samples were then analyzed via MALDI-TOF mass spectrometry and/or denaturing PAGE (20% acrylamide/bisacryamide, 75 W, 3 h). Gels were scanned on an Amersham Typhoon Biomolecular Imager and analyzed using ImageJ software.

*Phosphine screening*
The protocol in the general procedure above was followed using 5′-Cy5 labeled Mango II as the aptamer. All phosphines except for CyDPPDS were obtained from commercial sources. PTA, PTABS, TPP, TCEP, THPP, and THMP were purchased from Combi-blocks. CyDPP and DCyPDIBPS were obtained from Ambeed, and TPPTS was purchased from Sigma-Aldrich. Concentrated stocks were prepared by dissolving the phosphines in various ratios of DMSO and $H_2O$. PTA, PTABS, and TPPTS were dissolved in water. CyDPPDS, THMP, THPP, TCEP were dissolved in 20% DMSO in water. CyDPP, TPP, and DCyPDIBPS were dissolved in DMSO. Reactions were incubated at 37 °C for 2 h with shaking. Crosslinking was analyzed via gel electrophoresis as above.

*Ligand concentration dependence*
The protocol in the general procedure above was followed using 5′-Cy5 labeled Mango II as the aptamer. The RNA was incubated with increasing concentrations of **TO-1-CpO** (**1**) (5 –1000 µM) at 37 °C for 10 min with shaking. Reactions were treated with PTA (10 mM, $H_2O$) and incubated at 37 °C for 2 h. Crosslinking was assessed by denaturing PAGE as described above.

*Time dependence*
The protocol in the general procedure above was followed using 5′-Cy5 labeled Mango II as the aptamer. The only parameter that was varied was the reaction time (initial to 5 h). After crosslinking reactions were completed, samples were immediately analyzed by denaturing PAGE as described above.

*Temperature dependence*
The protocol in the general procedure above was followed using 5′-Cy5 labeled Mango II as the aptamer. 5′-Cy5 labeled Mango II (10 µM) was incubated with **TO-1-CpO** (**1**) (50–250 µM) at 4 °C, 20 °C, or 37 °C for 10 min with shaking. After pre-incubation with TO-1-CpO (**1**) at their respective reaction temperatures, the mixtures were treated with PTA (10 mM) and incubated at 4 °C, 20 °C, or 37 °C for 2 h with shaking. Results were assessed using denaturing PAGE.

*Competition experiments*
The protocol in the general procedure above was followed using 5′-Cy5 labeled Mango II as the aptamer. The RNA was treated with **TO-1-CpO** (**1**) (150 µM) in the presence of with increasing concentrations of TO-1-PEG$_3$-biotin (DMSO, 75–500 µM) to allow for competition before phosphine addition. Samples were incubated at 37 °C for 10 min with shaking followed by treatment with PTA (10 mM) at 37 °C for 2 h. Crosslinking was analyzed by denaturing PAGE as described above.

*Reactant specificity*
The protocol in the general procedure above was followed using 5′-Cy5 labeled Mango II as the aptamer. The RNA was incubated with various concentrations of **TO-1-CpO** (**1**) (0–250 µM), TO-1-NH$_2$ (**2**) (50–250 µM), or Ph-CpO (**3**) (50–250 µM) at 37 °C for 10 min while shaking. The reaction mixtures were then treated with PTA (10 mM, $H_2O$), and incubated at 37 °C for 2 h. Crosslinking was analyzed by denaturing PAGE as described above.

*Mutant Mango II crosslinking*
The protocol in the general procedure above was followed using 5′-Cy5 labeled Mango II as the wild-type aptamer. 5′-Cy5 labeled Mango II (10 µM), or mutant Cy5-Mango-II aptamer (10 µM) were prepared as before and incubated with **TO-1-CpO** (250 µM) for 10 min at 37 °C. Crosslinking was triggered by administration of PTA (10 mM) , and reactions were incubated for 2 h at 37 °C. Results were analyzed by denaturing PAGE as described above.

*Testing off target labeling with other nucleic acid motifs*
The protocol in the general procedure above was followed using various Cy5 labeled RNA samples (Mango-II, hTERT, c-Myc IRES WT D2 short, MycN, RNaseP, NRAS) (10 µM). RNAs were folded in 1X folding buffer (10 mM Tris (pH 7.0), 140 mM KCl, 1 mM $MgCl_2$) at 95 °C for 2 min followed by cooling on ice for 5 min. After folding, the RNA was incubated with either control (DMSO) or **TO-1-CpO** (**1**) (150 µM) at 37 °C for 10 min with shaking. Reactions were then treated with reactive phosphine PTA (10 mM, $H_2O$) at 37 °C for 2 h and analyzed by denaturing PAGE as described above. Samples were loaded at various time points onto the gel to ensure all samples would fit within one image. Larger RNA samples were loaded onto gel first and allowed to run before smaller RNA samples were loaded onto gel.

**Elucidation of crosslinking location**

*Purification of crosslinked RNA*
Crosslinking reaction was scaled up by 20-fold to a total volume of 200 µL following the general crosslinking protocol. Excess ligand and phosphine were filtered out using an Amicon Ultra centrifugal filter with a 3 kDa cutoff. Reaction was then buffered exchanged with the filter into the folding buffer. Crosslinked RNA was then separated using an AKTA Purifier connected to an anion exchange column (DNAPac PA-100, BioLC, 4 X 250 mm). Mobile phases used for purification were 1 mM Tris for solvent A and 1M KCl for solvent B. Both solvents were degassed prior to use. The method used for the purification was 5% B for 2.5 column volumes (CV), a 35% to 100% B

gradient over 6 CVs, 100% B for 1 CVs and then back to 5% B. RNA eluted from 17 to 26 mL. Fractions were collected in 1.6 mL aliquots. Each fraction was concentrated using the Amicon Ultra filter and buffer exchanged with TE buffer. To prepare for precipitation, each fraction was brought up to a final concentration of 300 mL of 300 mM KCl, and 1 µL of glycogen and 700 µL of ethanol were added. The mixture was stored in the –80 °C freezer overnight. The samples were spun at 16,000 rpm for 15 min to pellet the RNA, and then the supernatant was carefully removed. The pellet was washed with 400 mL of 70% ethanol and centrifuged again at the same setting. The supernatant was removed, and the pellet was dried under vacuum. Denaturing PAGE was then used to determine which fractions contained the crosslinked species.

*3′-End labeling of Mango with Cy5*
To a PCR tube, 1 µL of 10x T4 ligase buffer (New England Biolabs), 1 µL of 10 mM ATP, 1 µL of 100 mM of Mango II, 1 µL of DMSO, 1 µL of 100 mM of pCP-Cy4, 1 µL of T4 RNA ligase (New England Biolabs), and 4 µL of water were added and incubated at 16 °C overnight. EDTA was added at the end of the reaction to quench metal ions. Phenol-chloroform extraction was performed to extract the RNA. RNA was then precipitated as described previously and stored as a pellet in the -80 °C freezer until needed.

*RNAse T1 digestion*
For this assay, commercial 5′-Cy5 labeled Mango II and in-house made 3′-Cy5 labeled Mango II were both diluted to a final concentration of 10 µM. A pellet of an FPLC fraction that contained the crosslinked species was resuspended in 10 µL of TE buffer. To digest these RNAs, 1 µL of a 250 mM sodium citrate solution (pH 7.4) and 1 µL of 0.5 U/mL was added to 3 µL of each RNA. Digestion times ranged from 2 to 15 min at 60 °C, and reactions were quenched with 3 µL of 7 M urea, 40 mM EDTA, and orange G dye in 0.5x TBE. Denaturing PAGE was used to assess the results. Gel was first scanned for fluorescence and then stained with SYBR and imaged again.

*Reverse transcription (RT)*
RNA pellets were resuspended in TET buffer (10 mM Tris-HCl (pH 7.5), 0.1 mM EDTA, and 0.001% Triton X-100). Subsequently, 0.18 mM dNTPs, 0.02 mM biotinylated dUTP and dATP (Jena Bioscience) and 25 nM of the reverse primer were added to each reaction. The RNA and primer were annealed by heating at 90 °C for 1 min and cooled on ice for 3 min before the addition of 1X MMLV buffer (New England Biolabs), 200 U of MMLV reverse transcriptase (New England Biolabs) to each reaction. The reaction was initiated at 37 °C for 10 min and then the temperature was ramped up to 42 °C for 1 hr. Finally, the enzymes were inactivated at 75 °C for 5 min.

RT primer: 5′ -GTGTGCTCTTCCGATCT- 3′

*Binding to streptavidin beads*
5 U of RNaseH (New England Biolabs) was added to each RT reaction and incubated at 37 °C for 1 h. cDNA was then buffer-exchanged into G10 Sephadex beads with TET buffer by centrifugation at 2000 x*g* to remove free biotinylated dATP and dUTP. The cDNA was incubated with magnetic streptavidin beads (Dynabeads™ MyOne™ C1, Invitrogen) that were pre-equilibrated with streptavidin wash buffer (SWB; 60 mM Tris, 50 mM KCl and 0.001% Triton X-100) and incubated for 20 min at room temperature. After incubation, five washes were performed with SWB to remove excess RT primer.

*Addition of universal 5′ PCR adapter to 3′ of cDNA*
Ligation was performed on the magnetic beads following the RT step. The 3′ end of the cDNA was ligated with a 5′ phosphorylated splint hairpin containing a 3′ carbon spacer to attach a

constant-sequence region (T7 RNA polymerase A1 promoter sequence) to minimized sequences. The ligation was performed with a final concentration of 1X T4 DNA ligase reaction buffer (New England Biolabs) and 400 U of T4 DNA ligase (New England Biolabs), 1 mM rATP, 0.5 M betaine, 20% PEG8000 and 5 nM of splint hairpin. The reaction was incubated at 16 °C for 6 h, 30 °C for 6 h and inactivated at 65 °C for 15 min.

Splint hairpin for 1st round of *in vitro* selection: 5′ -P-
CCTATAGTGAGTCGTATTAAAAAAATATAGGNNNNNNN- 3′SpC3

The DNA oligos contained a 5′ phosphate to facilitate ligation by T4 DNA ligase and a 3′ carbon spacer to prevent self-ligation. Both oligos were predicted to fold into a hairpin secondary structure with a random sequence (NNNNNNN) acting as a non-sequence-specific splint.

The ligated cDNA was washed five times with SWB and incubated for 5 min on an orbital shaker to remove excess splint hairpin oligo. Two washes with 200 mM KOH were performed to remove cDNA from the magnetic streptavidin beads. After, elutions were pooled and precipitated with 1 µl of GlycoBlue (Invitrogen, MA, USA) and 2.5 volumes of cold 100% ethanol, and pelleted by centrifugation. The pellet was resuspended in TET buffer.

*Library amplification by PCR*

PCR amplification was performed on cDNA using a final concentration of a 1X Standard *Taq* reaction buffer (New England Biolabs), 0.2 mM dNTPs, 0.5 µM forward and reverse primer, and 1.25 U of DNA *Taq* polymerase (New England Biolabs). The reaction was amplified at 95 °C for 30 s, 50 °C for 30 s, and 72 °C for 30 s.

Forward primer (1st selection round): 5′ - CCTATATTTTTTAATACGACTCACTATAGG - 3′
Reverse primer: 5′ - GTGTGCTCTTCCGATCT - 3′

*HTS sequencing, processing, and mapping*

DNA was prepared for sequencing by first amplifying the DNA library with an Illumina forward adapter and reverse primer (above) to incorporate the Illumina adapter during PCR. The reaction was amplified for 4 to 8 cycles (95 °C for 30 s, 55 °C for 30 s, and 72 °C for 30 s). After amplification, the Illumina barcodes were added in a second PCR reaction and amplified for 1 cycle (95 °C for 30 s, 55 °C for 30 s, and 72 °C for 30 s) and then 7 cycles (95 °C for 30 s, 65 °C for 30 s, and 72 °C for 30 s). Both PCR reactions contained a final concentration of a 1X Standard *Taq* reaction buffer (New England Biolabs), 0.2 mM dNTPs, 0.5 µM forward and reverse primer, and 1.25 U of DNA *Taq* polymerase (New England Biolabs). All PCR products were visualized on a 3% agarose gel prior to purification with a DNA Clean and Concentrator kit (Zymo Research). The DNA libraries were then submitted for Illumina sequencing.

Illumina forward adapter: 5′ -
ACGACGCTCTTCCGATCTCCTATATTTTTTAATACGACTCACTATAGG- 3′

Sequenced reads were obtained using an Illumina MiSeq. Bowtie2 was used to map reads to the Mango II aptamer. The parameters -N 1 and --local were used when aligning the aptamer to their respective sequences. Bedtools genomecov was used to determine the most common sites where TO-1-CpO was reactive with the RNA.

**Oligonucleotides**

Deprotected and HPLC purified aptamers were purchased from Horizon Discovery Inc. or IDT with the following sequences and used without any further purifications.

| Oligo | Sequence |
|---|---|
| Mango I | GGCACGUACGAAGGGACGGUGCGGAGAGGAGAGUACGUGC |
| Mango II | GGCACGUACGAAGGAGAGGAGAGGAAGAGGAGAGUACGUGC |
| Mango III | GGCACGUACGAAGGAAGGAUUGGUAUGUGGUAUAUUCGUACGUGCC |
| Mango IV | GGCACGUACCGAGGGAGUGGUGAGGAUGAGGCGAGUACGUGC |
| Mango II mutant | CCCACGUACGAAGGAGACCACAGGAAGAGGAGAGUACGUGC |
| Cy5-Mango I | Cy5- GGCACGUACGAAGGGACGGUGCGGAGAGGAGAGUACGUGC |
| Cy5-Mango II | Cy5- GGCACGUACGAAGGAGAGGAGAGGAAGAGGAGAGUACGUGC |
| Cy5-Mango III | Cy5- GGCACGUACGAAGGAAGGAUUGGUAUGUGGUAUAUUCGUACGUGCC |
| Cy5-Mango IV | Cy5- GGCACGUACCGAGGGAGUGGUGAGGAUGAGGCGAGUACGUGC |
| Cy5-Mango II mutant | Cy5-CCCACGUACGAAGGAGACCACAGGAAGAGGAGAGUACGUGC |
| Cy5-Mango II primer | Cy5-GGCACGUACGAAGGAGAGGAGAGGAAGAGGAGAGUACGUGCAGAUCGGAAGAGCACAC |
| Cy5-MyCN | Cy5-AGGGGGTGGGAGGGGGCATGCAGATGCAGGGGGT |
| Cy5-RNaseP | TGTGTAAGTTTTAGGTTGATTTG-Cy5 |
| Cy5-NRAS | Cy5-UGUGGGAGGGGCGGGUCUGGG |
| Cy5-hTERT | Cy5-AGGGGGCTGGGCCGGGGACCCGGGAGGGGTCGGGACGGGGCGGGGT |
| Cy5-cMyc IRES WT D2 short | Cy5-GCGGGGAGGCUAUUCUGCCCAUUUGGGGACACUUCCCCGCCGCUG |

### General synthetic procedures

Unless otherwise stated, all reagents and solvents were purchased from commercial vendors and used as received. CpO-PFP[1], CyDPPDS[2] and TO-1-acetate[3] were synthesized following literature precedent. Anhydrous organic solvents were prepared by degassing with argon and passing through two 4 x 36 in. columns of anhydrous neutral A2 (8 x 12 mesh; LaRoche Chemicals; activated at 350 °C for 12 h under a flow of argon). Column chromatography was carried out using Silicycle 60 Å (32–64 mesh) silica gel. Reactions were monitored by TLC and LC-MS. Thin layer chromatography (TLC) was carried out with Merck Millipore 250 mm silica gel F-254 plates, and plates were visualized using UV light or KMnO4 stain. Organic solutions were concentrated under reduced pressure using a Büchi rotary evaporator. A CombiFlash Rf purification system with RediSep Silver flash silica gel or $C_{18}$ columns were used for some column chromatography experiments. HPLC purifications were performed on an Agilent 1260 Infinity II equipped with UV-Vis Detector, using an Agilent Eclipse XDB-C18column (5 mm, 9.4 x 250 mm) with a 4 mL/min flow rate. Mobile phases used for the HPLC were water with 0.1% formic acid for solvent A and acetonitrile with 0.1% formic acid for solvent B. HPLC runs were monitored at $\lambda$ = 254 and 500 nm.

All reported $^1$H-NMR spectra were measured at 500 MHz, and the complementary $^{13}$C NMR frequency was 126 MHz. Reported $^1$H NMR chemical shifts are in parts per million with the solvent resonance as an internal standard ($CD_3OD$: 3.31 ppm). Reported $^{13}$C NMR chemical shifts are in parts per million with the solvent resonance as the internal standard ($CD_3OD$: 49.0 ppm). NMR data are processed, annotated, and reported in the following format: chemical shift (multiplicity (s = singlet, br s = broad singlet, d = doublet, t = triplet, q = quartet, m = multiplet), coupling constants (Hz), Integration) using MestReNova. High-resolution mass spectrometry (HRMS) data were obtained with an LC–MS time-of-flight mass spectrometer by electrospray ionization (ESI). Mass spectra were acquired at the University of California, Irvine Mass Spectrometry Facility and NCI Biophysics Resource Center.

### Synthetic procedures

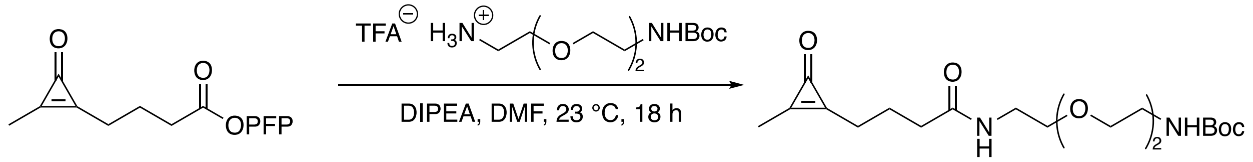


CpO-PFP

**CpO-PEG$_2$-NHBoc (1)**

**CpO-PEG$_2$-NHBoc (1)**. To a flame-dried round bottom flask was added *N*-Boc-2,2′-(ethylenedioxy)diethylamine (0.112 g, 0.450 mmol, 1.2 equiv.), DIPEA (0.135 mL, 0.375 mmol, 2.0 equiv.), and anhydrous DMF (3.5 mL). CpO-PFP[1] (0.120 g, 0.375 mmol, 1.0 equiv.) was added as a solid, and the resulting solution was stirred at room temperature overnight under nitrogen. The reaction was then concentrated *in vacuo*, and the crude reaction mixture was purified by flash chromatography (2–10% MeOH in $CH_2Cl_2$) to give **1** as a yellow oil (0.133 g, 92%). $^1$H NMR (400 MHz, $CD_3OD$) δ 3.61 (s, 4H), 3.57 (t, *J* = 5.3 Hz, 4H), 3.47 (q, *J* = 5.4 Hz, 2H), 3.32 (s, 2H), 2.67 (t, *J* = 7.0 Hz, 2H), 2.37 (t, *J* = 7.4 Hz, 2H), 2.27 (s, 3H), 2.04 (q, *J* = 7.1, 2H), 1.45 (s, 9H). $^{13}$C NMR (151 MHz, $CDCl_3$) δ 171.9, 160.9, 160.1, 157.0, 156.1, 79.3, 70.3, 70.2, 69.8, 40.4, 39.2, 35.0, 35.0, 28.4, 25.4, 22.1, 11.3. HRMS (ESI+) calculated for $C_{19}H_{32}N_2O_6Na$ $[M+Na]^+$ *m*/*z*: 407.2158; found 407.2144.

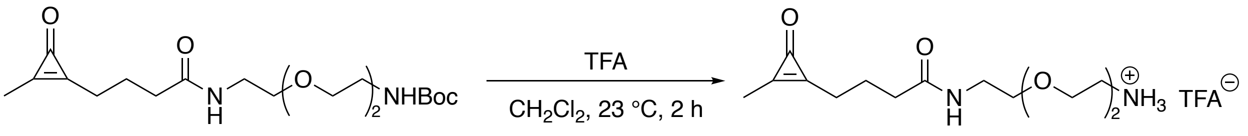


**CpO-PEG$_2$-NHBoc (1)** **CpO-NH$_2$ (2)**

**CpO-NH$_2$ (2)**. To a scintillation vial containing compound **1** (0.133 g, 0.327 mmol) was added $CH_2Cl_2$ (2 mL). TFA (2 mL) was added dropwise, the vial was sealed, and the solution was stirred at room temperature for 2 h. The crude reaction mixture was concentrated *in vacuo*, redissolved in MeOH (1 mL), and precipitated with cold diethylether (45 mL). The product suspension was filtered and washed with cold diethylether (2 x 45 mL) to give **2** as a tacky, yellow solid (0.086 g, 87%). $^1$H NMR (400 MHz, $CD_3OD$) δ 3.68 (t, *J* = 5.1 Hz, 2H), 3.64 (m, 5H), 3.54 (t, *J* = 5.5 Hz, 2H), 3.36 (t, *J* = 5.8Hz, 2H), 3.11 (t, *J* = 4.8Hz, 2H), 2.68 (t, *J* =7.0Hz, 2H), 2.32 (t, *J* = 7.3Hz, 2H), 2.29 (s, 3H) 2.00 (q, *J* = 7.3 Hz, 2H). $^{13}$C NMR (151 MHz, $CD_3OD$) δ 70.0, 69.9, 69.2, 39.3, 38.8, 34.6, 24.6, 21.9, 9.1. HRMS (ESI+) calculated for $C_{14}H_{25}N_2O_4$ $[M+H]^+$ *m*/*z*: 285.1814; found 285.1805.

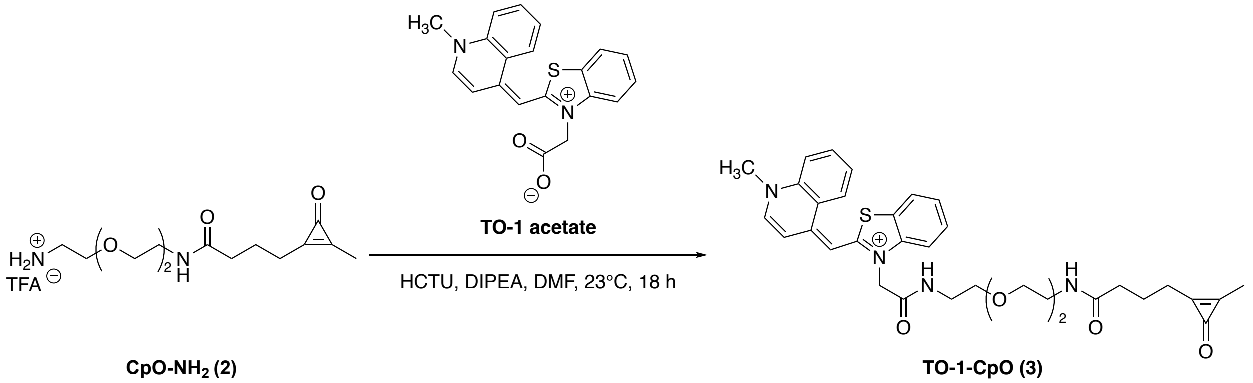


**CpO-NH$_2$ (2)** **TO-1-CpO (3)**

**TO-1-CpO (3)**. TO-1-acetate (0.057 g, 0.16 mmol, 0.9 equiv.), DIPEA (0.047 g, 0.36 mmol, 2,0 equiv.), and HCTU (0.15 g, 0.36 mmol, 2.0 equiv.) were added to a flame-dried round bottom flask containing DMF (4 mL). The solution was stirred for 10 min. **2** (0.052 mg, 0.18 mmol, 1.0 equiv.) was then added, and the reaction was stirred at room temperature for 18 h. DMF was removed via toluene azeotrope, and the sample was HPLC purified (isocratic method at 22% B for 16 min). The product eluted at 11.4 min and was collected. Another HPLC purification was performed using a 10-90% gradient of B over 16 min. The desired fraction at $r_T$ ~7.5 min was collected. The collected fractions were lyophilized, yielding a red powder (0.008 g, 8%). Compound purity was verified using HPLC (eluting with 10-90% solvent B over 16 min, Figure S9). HRMS (ESI+) calculated for $C_{34}H_{39}N_4O_5S^+$[M+] *m*/*z*: 615.2641; found 615.2627.

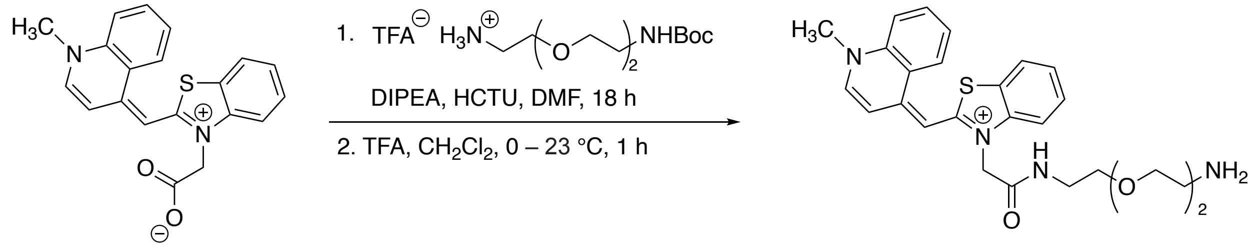


**TO-1 acetate** **TO-1-NH$_2$ (4)**

**TO-1-NH$_2$ (4)**. To a flame-dried round bottom flask, 2 mL of DMF, TO-1 acetate (0.053 g, 0.15 mmol, 1.1 equiv.), followed by HCTU (0.062 g, 1.9 mmol, 1.1 equiv.) and DIPEA (0.071 g, 3.8 mmol, 4.0 equiv.) were added under nitrogen gas. The reaction vessel was covered in aluminum foil, and the reaction was stirred for 5 min at room temperature. Next, *N*-Boc-2,2′-(ethylenedioxy)diethylamine (0.034 g, 0.14 mmol, 1.0 equiv) was added, and the reaction was stirred at room temperature overnight. The reaction was concentrated in vacuo, and resulting crude was purified by silica gel chromatography to yield TO-1-NHBoc (24 mg, 31% yield) of the intermediate product. Next, the purified intermediate was then dissolved in DCM (3.6 μL), put under a nitrogen atmosphere, and cooled to 0 ºC. After, 0.50 mL (excess) of trifluoroacetic acid (TFA) was added to the reaction mixture dropwise. The reaction was allowed to warm to room temperature and stirred for 1 hour. The reaction progress was monitored by LCMS and upon completion was purified via reverse phase $C_{18}$ flash chromatography to yield the final product: 14 mg, 42%, red solid. $^1$H NMR (500 MHz, $CD_3OD$) δ 8.56-8.42 (m, 2H), 8.07-7.96 (m, 2H), 7.89-7.74 (m, 2H), 7.59-7.48 (m, 3H), 7.43-7.36 (m, 1H), 5.28 (s, 2H), 4.20 (s, 3H), 3.66-3.51 (m, 10H), 3.04 (t, $J$ = 5.2 Hz, 2H), 1.39-1.35 (m, 2H); $^{13}$C NMR (126 MHz, $CD_3OD$) δ 168.0, 162.0, 151.2, 146.1, 141.9, 139.8, 134.7, 129.4, 128.5, 126.2, 125.9, 125.9, 125.5, 123.7, 119.1, 113.3, 110.2, 71.3, 71.3, 70.6, 67.9, 55.8, 43.3, 40.6, 18.7, 17.3, 13.1. HRMS (ESI+) calculated for $C_{26}H_{31}N_4O_3S^+$ $[M]^+$ *m*/z: 479.2111; found 479.2121.

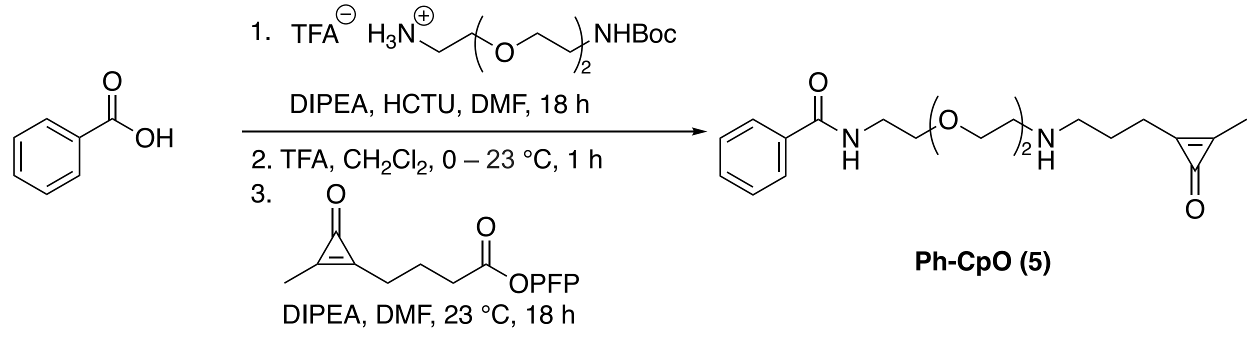


**Ph-CpO (5)**

**Ph-CpO (5).** To a flame-dried round bottom, 3 mL of DMF, benzoic acid (0.081 g, 0.66 mmol, 1.1 equiv.), 275 mg (1.1 equiv.) HCTU (0.28 g, 0.66 mmol, 1.1 equiv.), and diisopropylethylamine (0.63 g, 4.8 mmol, 8.0 equiv.) were added to a reaction vessel under an atomosphere of nitrogen gas, and stirred for 5 minutes at room temperature. Next, *N*-Boc-2,2′-(ethylenedioxy)diethylamine (0.15 g, 0.60 mmol, 1.0 equiv.) was added to the reaction vessel and allowed to stir at room temperature overnight. The reaction was purified by silica gel chromatography to yield the Ph-PEG$_2$-NHBoc. Next, the purified intermediate was then dissolved in 5 mL of DCM, put under a nitrogen atmosphere, and cooled to 0 ºC. After, 1.0 mL (20.0 equiv.) of trifluoroacetic acid (TFA) was added to the reaction mixture dropwise, allowed to warm to room temperature and stirred for 1 hour. The reaction was purified via reverse phase $C_{18}$ flash chromatography to yield Ph-PEG$_2$-NH$_2$ (0.11 g, 52% yield). Finally, Ph-PEG$_2$-NH$_2$ (0.11 g, 0.31 mmol, 2.0 equiv.), was dissolved in 1 mL DMF under nitrogen gas, and DIPEA (0.12 g, 0.94 mmol, 6.0 equiv.), and CpO-PFP (0.050 g, 0.16 mmol, 1.0 equiv.) were sequentially added and stirred at room temperature overnight. The reaction was purified by HPLC to yield the final product: 41 mg, 68%, clear oil. $^1$H NMR (500 MHz, $CD_3OD$) δ 7.84-7.80 (m, 2H), 7.53 (t, *J* = 7.4 Hz, 1H), 7.46 (t, *J* = 7.7 Hz, 2H), 3.69-3.61 (m, 6H), 3.58 (t, *J* = 6.0 Hz, 2H), 3.54 (t, *J* = 6.0 Hz, 2H), 3.34 (t, *J* = 6.0 Hz, 2H), 2.66 (t, *J* = 7.1 Hz, 2H), 2.32-2.26 (m, 5H), 2.03-1.96 (m, 2H); $^{13}$C NMR (126 MHz, $CD_3OD$) δ 174.9, 170.3, 162.4, 161.6, 158.7, 135.7, 132.7, 129.6, 128.3, 71.3, 71.3, 70.6, 70.6, 40.9, 40.3, 35.9, 26.0, 23.3, 10.5. HRMS (ESI+) calculated for $C_{21}H_{28}N_2O_5$ $[M+H]^+$ *m*/z: 389.2071; found 389.2078.

### NMR Spectra

$^1$H NMR spectrum for **CpO-PEG$_2$-NH$_2$ (1)**

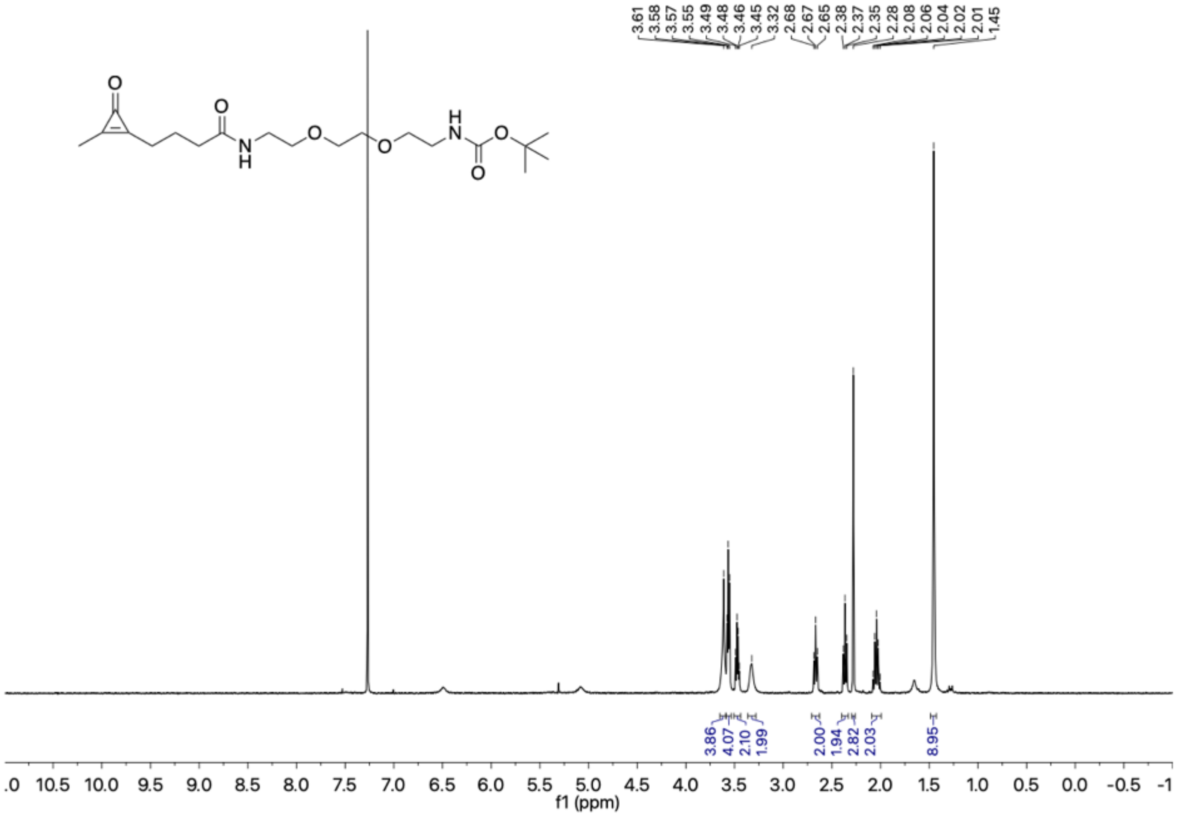

$^{13}$C NMR spectrum **CpO-PEG$_2$-NH$_2$ (1)**

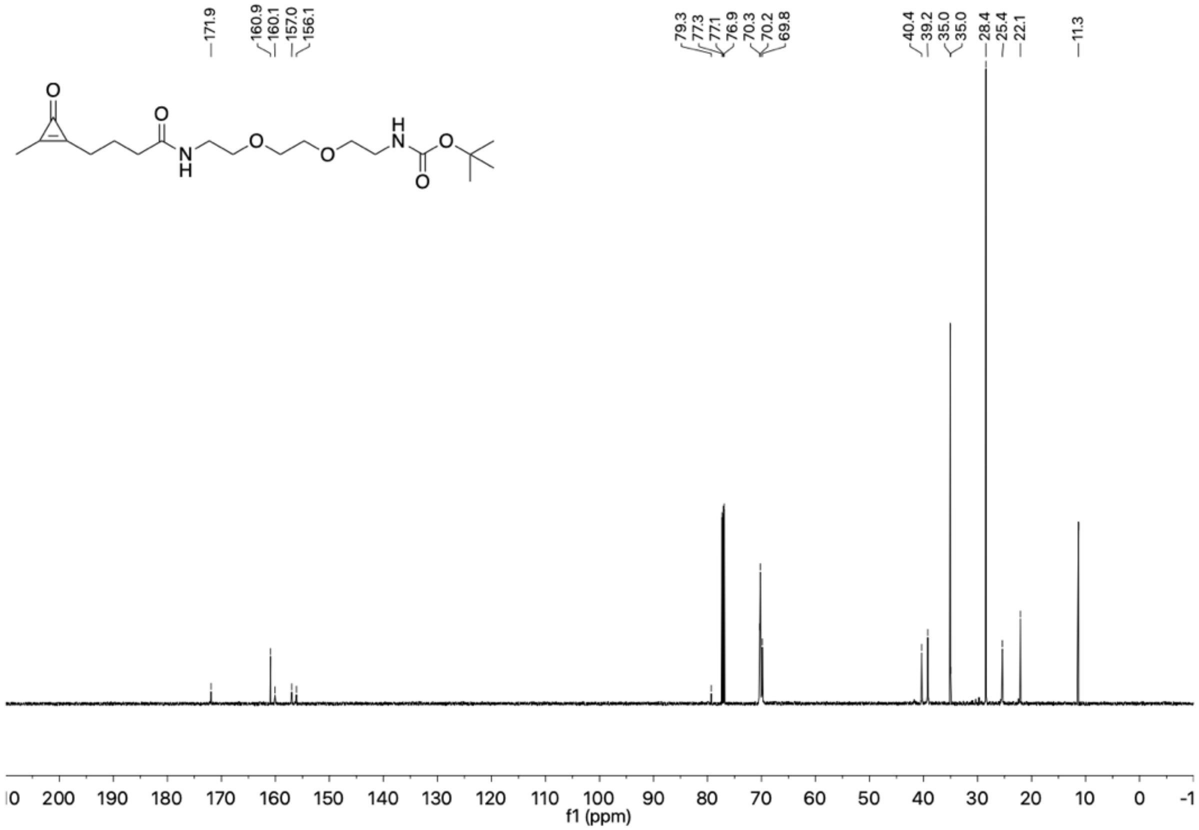

$^{1}$H NMR spectrum for **CpO-NH$_2$ (2)**

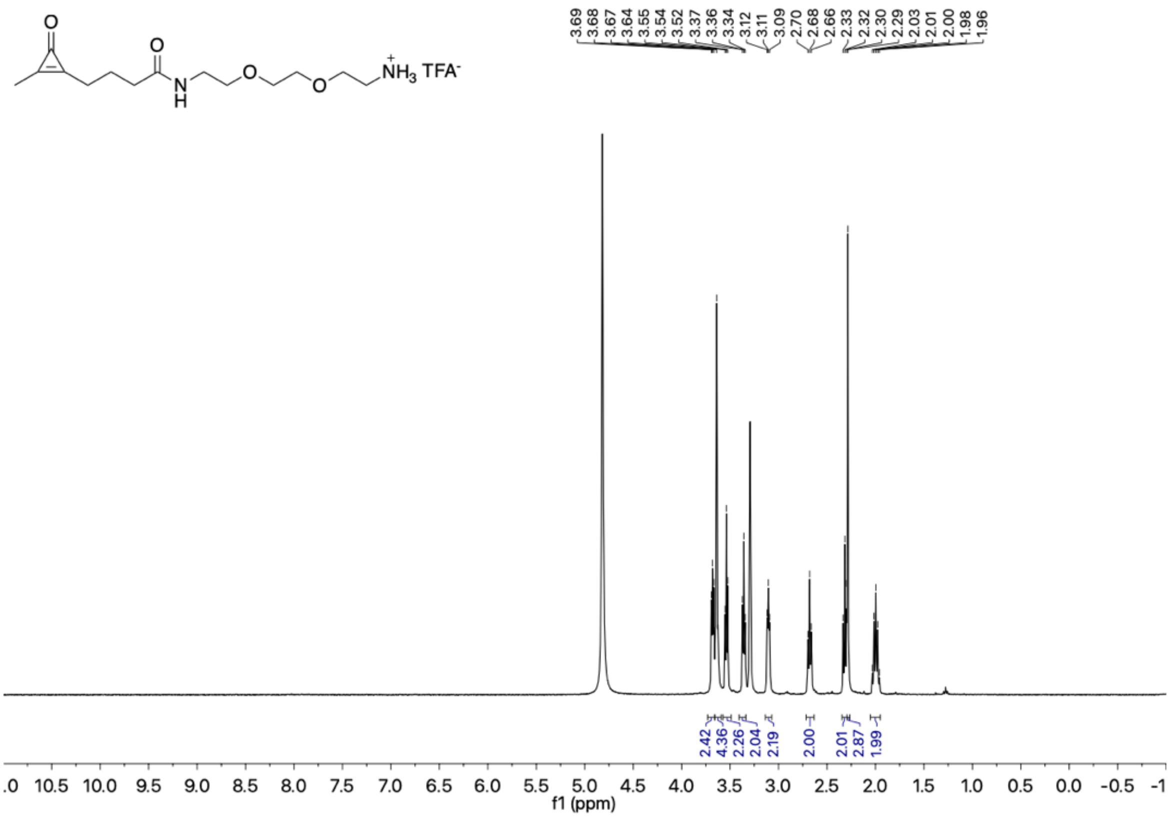

$^{13}$C NMR spectrum for **CpO-NH$_2$ (2)**

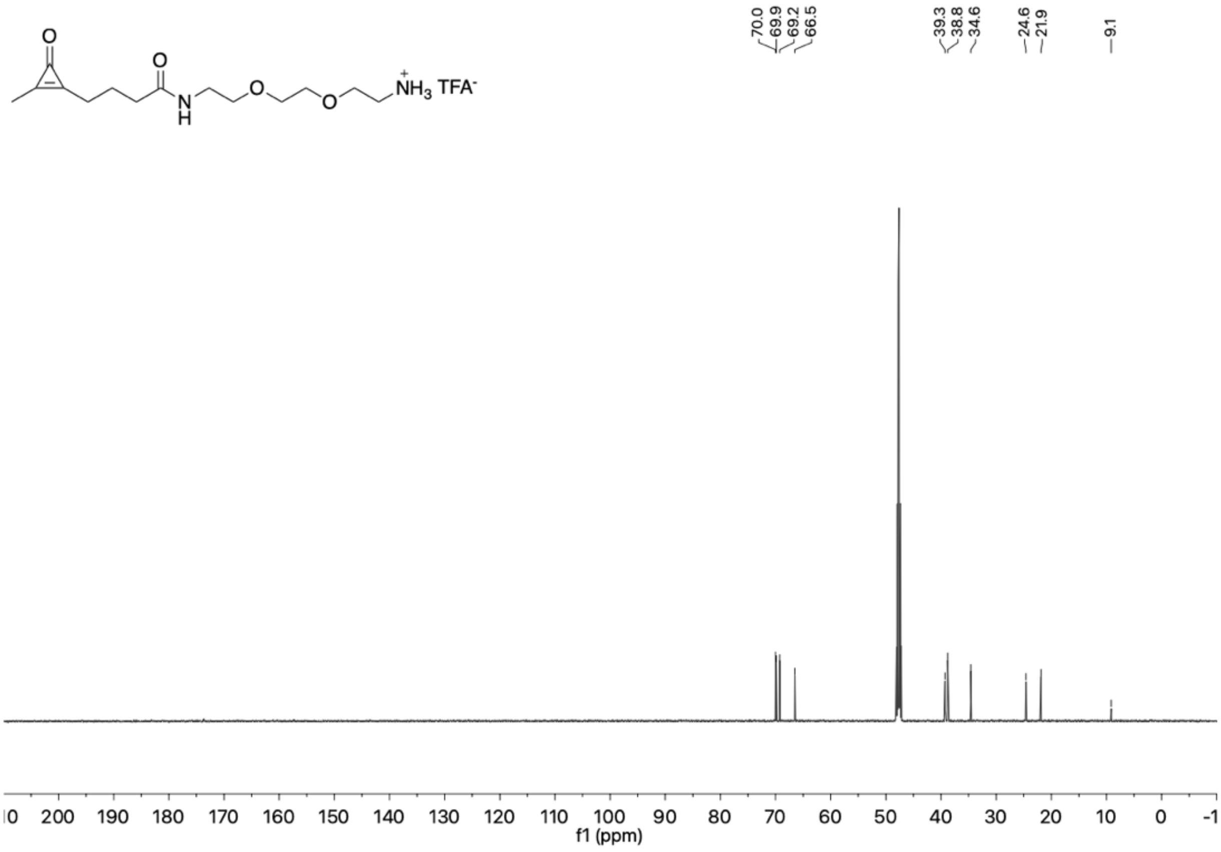

$^{1}$H NMR spectrum for **TO-1-NH$_2$** (**4**).

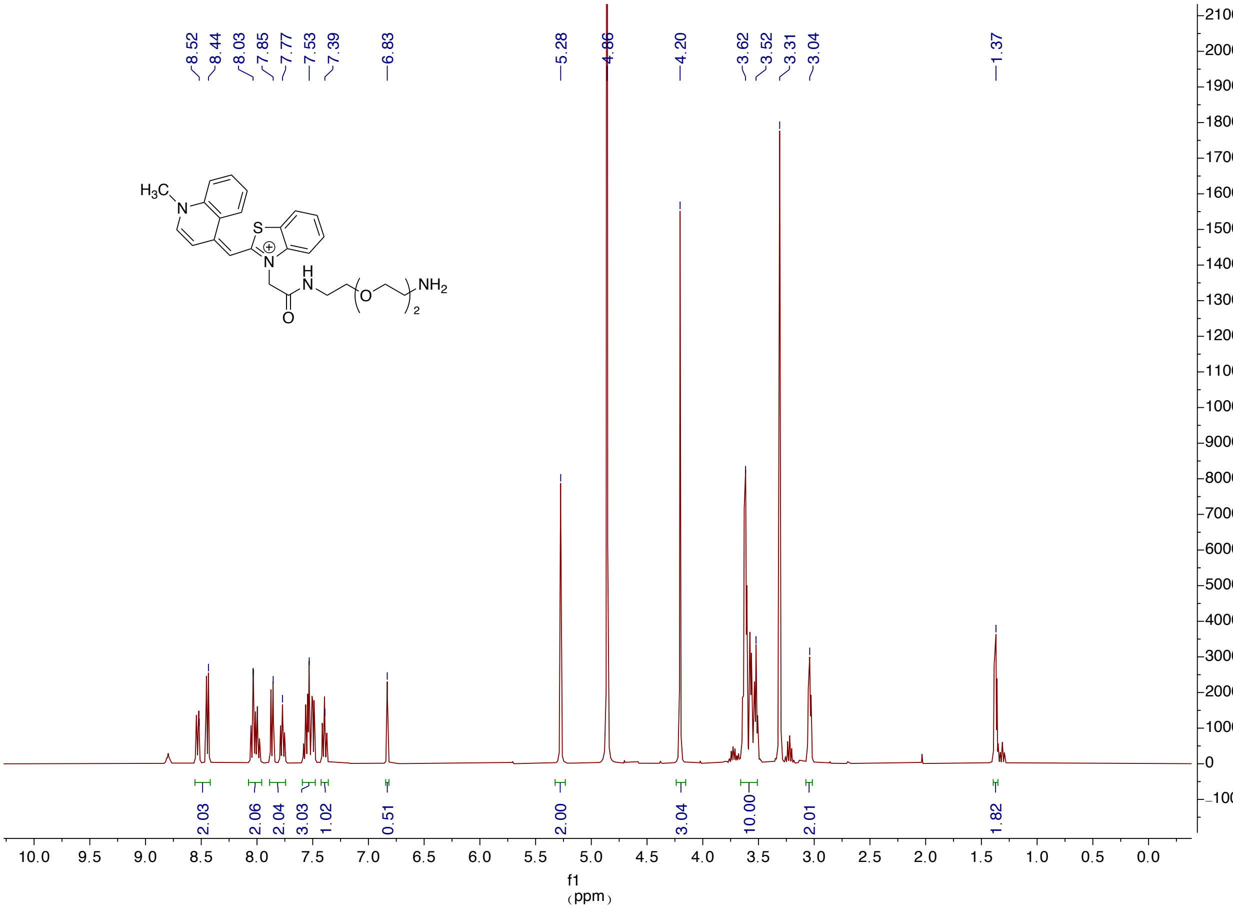

$^{13}C$ NMR spectrum for **TO-1-NH$_2$ (4)**.

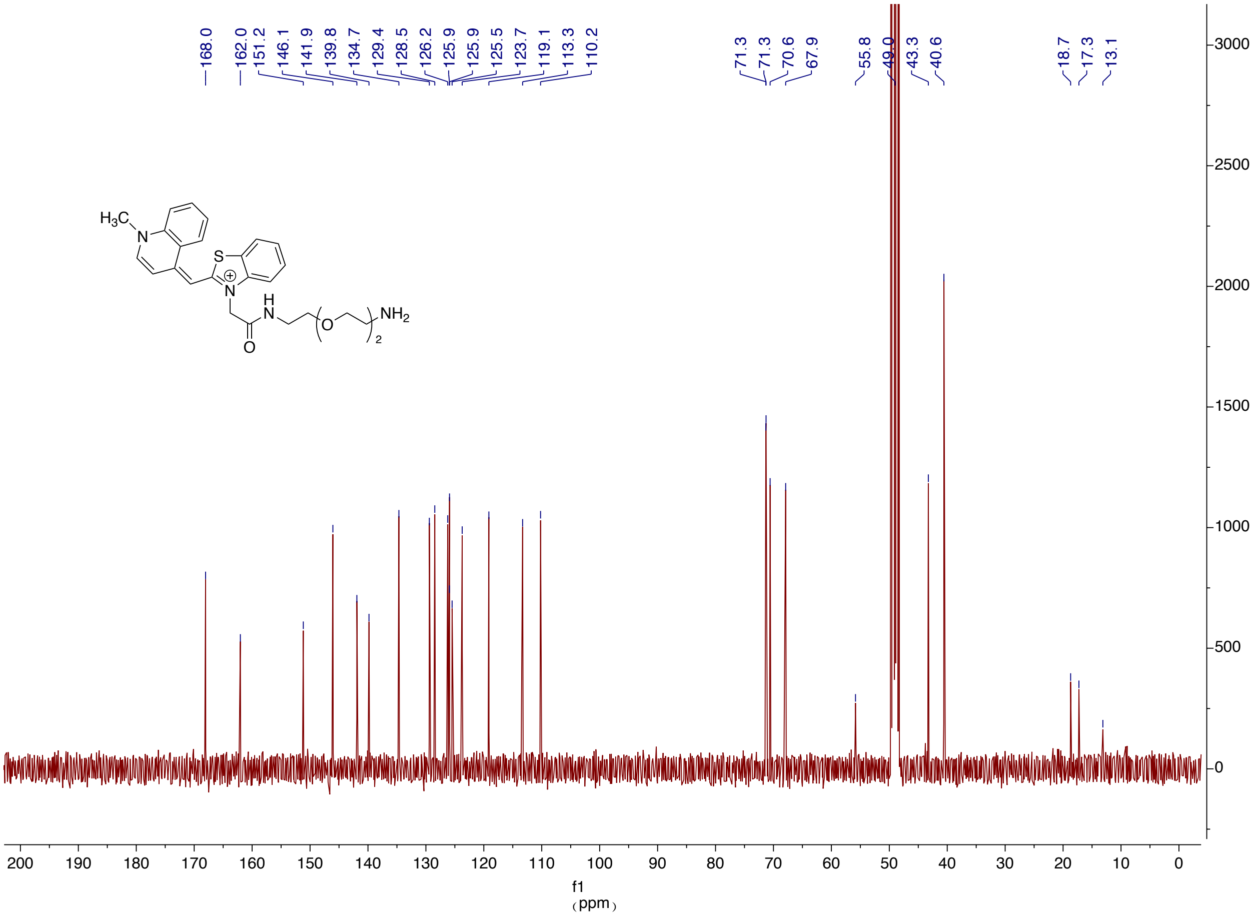

$^{1}$H NMR spectrum for **Ph-CpO** (**5**).

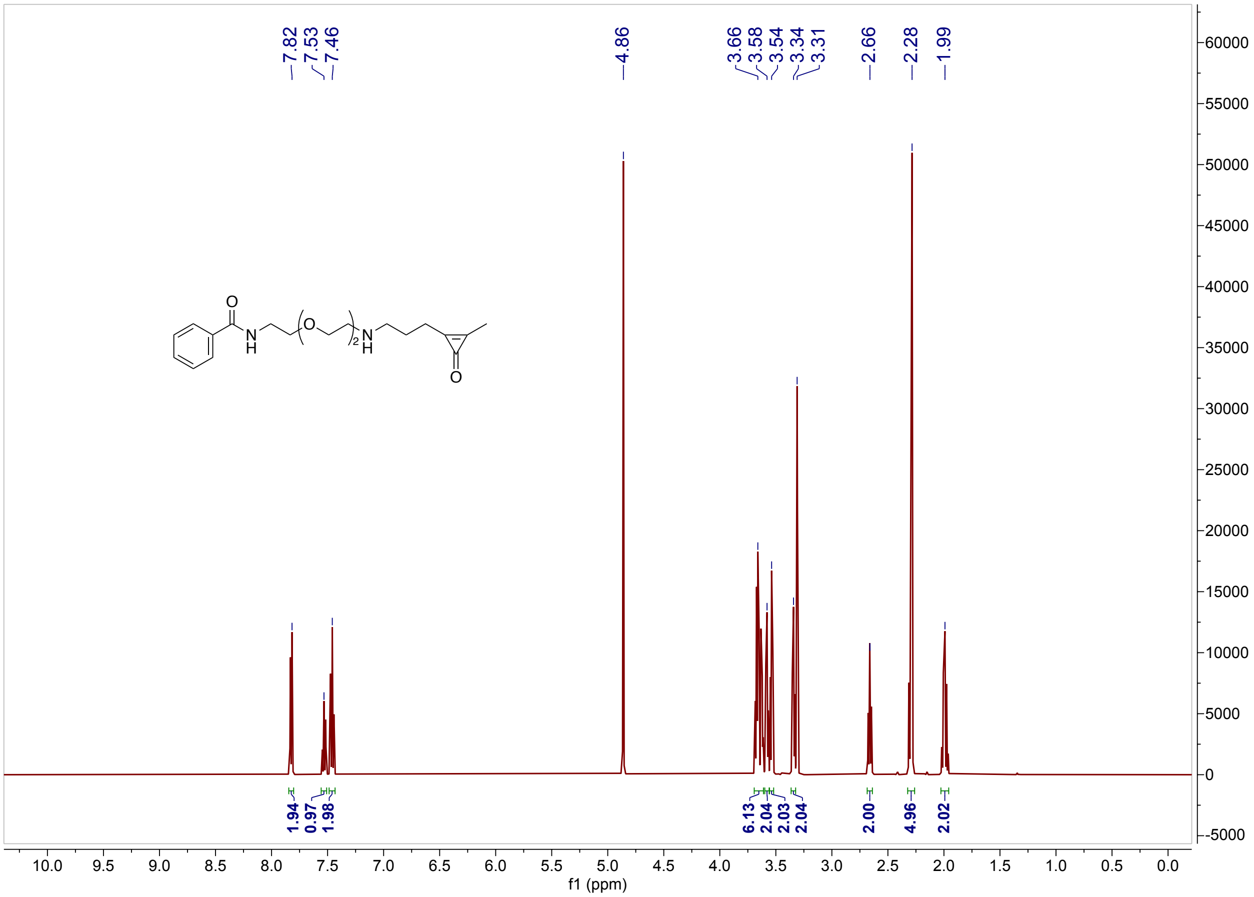

$^{13}$C NMR spectrum for **Ph-CpO** (**5**).

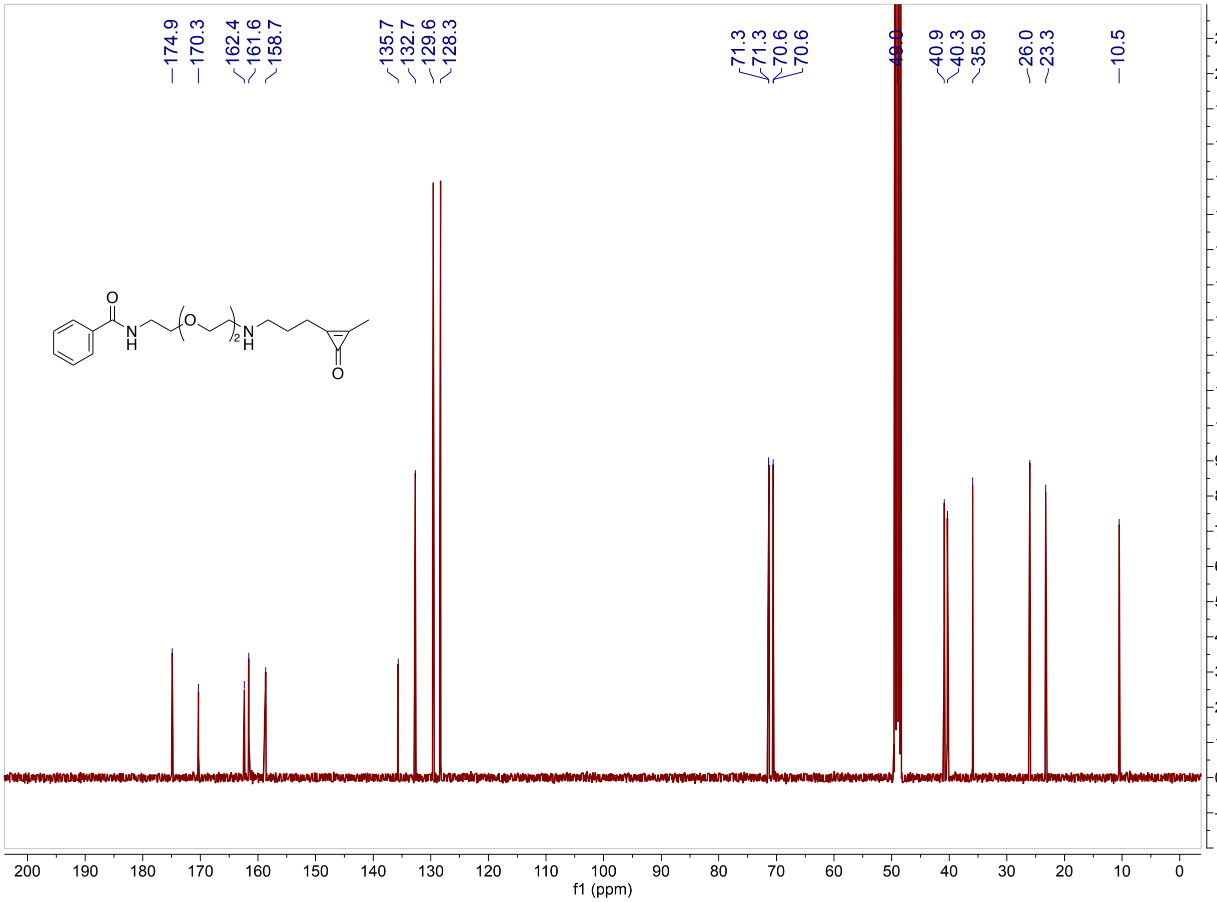
